# First come, first served: Superinfection exclusion in Deformed wing virus is dependent upon sequence identity and not the order of virus acquisition

**DOI:** 10.1101/2021.03.22.436467

**Authors:** Olesya N Gusachenko, Luke Woodford, Katharin Balbirnie-Cumming, David J Evans

**Affiliations:** Biomedical Sciences Research Complex, University of St. Andrews, North Haugh, St. Andrews, UK

**Author notes:** corresponding author, address: University of St Andrews, North Haugh, St Andrews, KY16 9ST UK.

## Abstract

Deformed wing virus (DWV) is the most important globally distributed pathogen of honey bees and, when vectored by the ectoparasite *Varroa destructor*, is associated with high levels of colony losses. Divergent DWV types may differ in their pathogenicity and are reported to exhibit superinfection exclusion upon sequential infections, an inevitability in a *Varroa*-infested colony. We used a reverse genetic approach to investigate competition and interactions between genetically distinct or related virus strains, analysing viral load over time, tissue distribution with reporter gene-expressing viruses and recombination between virus variants. Transient competition occurred irrespective of the order of virus acquisition, indicating no directionality or dominance. Over longer periods, the ability to compete with a pre-existing infection correlated with the genetic divergence of the inoculae. Genetic recombination was observed throughout the DWV genome with recombinants accounting for ~2% of the population as determined by deep sequencing. We propose that superinfection exclusion, if it occurs at all, is a consequence of a cross-reactive RNAi response to the viruses involved, explaining the lack of dominance of one virus type over another. A better understanding of the consequences of dual- and superinfection will inform development of cross-protective honey bee vaccines and landscape-scale DWV transmission and evolution.

## Introduction

Honey bees (*Apis mellifera*) are globally important pollinators of wild flowers and agricultural crops, and the source of honey, with annual global production worth in excess of $7bn [1]. Both honey production and pollination services require strong, healthy colonies, which are threatened by a range of factors, but most significantly by disease. One of the major viral pathogens of honey bees is Deformed wing virus (DWV). When transmitted by the parasitic mite *Varroa destructor,* DWV is responsible for high overwinter colony losses, which can exceed 37% annually [2]. Improvements to honey bee health, through direct control of virus transmission or replication, require a better understanding of how the virus propagates within and between bees.

The historical identification and naming of DWV-like viruses imply a greater genetic divergence than subsequent molecular analysis has demonstrated. In 2004-2006 several picorna-like viruses with high levels of sequence identity were reported [3–5]. These viruses were initially named according to their origins; the virus from honey bees with characteristic wing deformities was termed DWV [4], a similar virus found in aggressive workers in Japan was designated Kakugo virus [3, 5] and analysis of *Varroa* mites yielded Varroa destructor virus type 1 (VDV-1) [3]. Limited genetic divergence (~84-97% genomic RNA identity), similar infectivity in honey bees, and demonstrated ability to freely recombine during coinfections [6–9] resulted in them now being considered as different variants of DWV [6, 7, 10], albeit occupying two genetic branches (VDV-1-like and DWV-like) of the same phylogenetic tree [11]. To distinguish between these branches the terminology ‘type A’ and ‘type B’ has been adopted for DWV-like and VDV-1-like variants respectively. Evidence for the existence of a third type named DWV type C has also been reported [12].

DWV is ubiquitous in honey bees [13–15], with the possible exception of Australian colonies [16]. In the absence of *Varroa* the virus is transmitted horizontally, *per os*, and vertically from the infected queen and the drones [17]. With subsequent *Varroa* mite transmission it is therefore inevitable that the virus enters a host already harbouring one or multiple DWV variants. Current studies suggest that DWV infection can occur with several variants cocirculating in the same apiary, colony or individual honey bee host [18–21]. Although the type A and B variants appear to be differentially distributed, with type A frequently reported in the US and type B being commonly detected in European colonies [8, 13, 22], direct competition may occur where they cocirculate. If this competition has directionality it will influence the distribution and future spread of DWV at the landscape scale. While some studies of mixed DWV infections demonstrate no predominance of one variant over another [18, 23], others show possible competition between the variants and higher accumulation of DWV B in infected bees [24]. In addition, superinfection exclusion (SIE) has been proposed, in which a pre-existing type B virus prevents the establishment of a type A infection at the colony level [10].

A recently developed reverse genetics (RG) system comprising a set of genetically tagged DWV variants and reporter gene-expressing viruses provides an opportunity to investigate coinfection kinetics and competition between DWV types [25]. Since SIE is a widely observed virological phenomenon [26–35], we extended these studies to assay dominance of one variant over another during sequential infection. Using reporter gene-expressing DWV we additionally investigated the influence of competition on tissue distribution of infection. We show that where competition is observed, manifest as reduced virus levels, it is reflected in reduced reporter gene expression at the cellular level. Notably we show that DWV accumulation during superinfection is influenced by the genetic identity between the viruses, rather than by a directionality of competition. Genetically divergent DWV variants (such as those representing type A and type B) exhibit transient competition, whilst viruses with greater identity (*e.g.* type A/B recombinants with either type A or type B) demonstrate distinctly more pronounced effect. We also analysed the occurrence and identity of recombinants during mixed infections and confirmed that these are present with junctions widely distributed throughout the genome. These studies provide further insights into the biology of DWV. In particular they address the consequences of co- and superinfection, an important consideration when transmitted by the ectoparasite *Varroa*. Our results indicate that genome identity is the determinant that defines the outcome of dual infections; this will inform studies of population transmission at the landscape scale and possible future developments of ‘vaccines’ to protect honey bees from viral disease [36].

## Materials and Methods

### RG DWV clones preparation

VDD, VVD and VVV RG constructs used in this study were described earlier [25], DDD RG cDNA was prepared by modification of the VDD RG system with DWV type A parental sequence insert, which was based on published data [37] and obtained by custom gene synthesis (IDT, Leuven, Belgium). EGFP and mCherry-expressing chimeric DWV genomes were built via incorporation of the reporter-encoding sequence into DWV cDNA as described previously [25]. All plasmid sequences were verified by Sanger sequencing. cDNA sequences of DDD and VVV_mC_ are shown in Text S1, other RG cDNAs are available online (GenBank accession numbers: DWV-VDD - MT415949, DWV-VVD - MT415950, DWV-VVV - MT415952, DWV-VDD-eGFP - MT415948, DWV-VVD-eGFP - MT415953).

### Viral RNA and siRNA synthesis

DWV RNA was synthesized from linearized plasmid templates with T7 RiboMAX Express Large Scale RNA Production System (Promega, Southampton, UK), and purified with GeneJet RNA Purification Kit (Thermo Fisher Scientific) as described in [25].

siRNA strands were prepared using Express Large Scale RNA Production System (Promega) according to the manufacturer’s protocol with double stranded DNA templates annealed from synthetic oligonucleotide pairs containing T7 RNA polymerase promoter sequence (Table S1).

### Viruses

Infectious DWV was prepared from honey bee pupae injected with *in vitro* generated RNA as previously described [25, 38]. For quantification RNA was extracted from 100 μl of virus preparation using RNeasy kit (Qiagen, Manchester, UK) and analysed by reverse transcription and quantitative PCR (qPCR).

### Honey bees and bumble bees

All honey bee (*Apis mellifera*) brood in this study was obtained from the University of St Andrews research apiary. Colonies were managed to reduce *Varroa* levels and endogenous DWV levels were regularly tested. Honey bee larvae and both honey and bumble bee pupae (*Bombus terrestris audax*, Biobest, Belgium) were maintained and fed as described previously [25].

### Virus inoculations

Virus injections of pupae were performed with insulin syringes (BD Micro Fine Plus, 1 ml, 30 G, Becton Dickinson, Oxford, UK) as described in [25, 38].

Oral larval infection was carried out by single DWV feeding according to the previously described procedure [25]

### RNA extraction, reverse transcription and PCR (RT-PCR)

RT-PCR and qPCR analysis of individual pupae samples was performed as previously described [25]. Sequences of primers are shown in Table S1. When required, PCR products were subjected to restriction digest prior to loading on the 1% agarose gel stained with ethidium bromide. DWV titres were calculated by relating the resulting Ct value to the standard curve generated from a serial dilution of the cDNA obtained from the viral RNA used for virus stock preparation.

### Microscopy

Imaging was conducted using a Leica TCS SP8 confocal microscope with 10× HC PL FLUOTAR objective. For dissected pupae analysis samples were mounted in a drop of PBS under the microscope cover slides and observed by microscopy within 1 h after the dissection.

### Sample libraries for next generation sequencing

RNA was reverse transcribed using Superscript III polymerase (Invitrogen, Thermo Fisher Scientific) with DWV FG RP1 primer (Table S1) using 1 μg of total RNA in a 20 μl final reaction volume and following the manufacturer’s protocol. Reactions were incubated at 50°C for 1 h, 75°C for 15 min.

The transcribed cDNA was amplified using LongAmp Taq polymerase (New England Biolabs) to produce a ~10 Kb PCR fragment. The reactions were carried out according to the manufacturer’s protocol with the following thermal profile: 30 s at 95°C, 30 cycles of 95°C for 15 s, 53°C for 30 s and 65°C for 8 min, with a final extension at 65°C for 10 min.

### Recombination Analysis

Purified amplicons were sequenced using an Illumina Hi-seq at the University of St Andrews, producing 2×300 bp paired-end reads. The sequences were converted to Fasta format, extracted and trimmed using Geneious (v.2019.1.3). A reference genome file was made using VVV and VDD cDNA sequences with a terminal pad of A-tails added to maximise sensitivity [39]. The reference file was indexed using Bowtie Build (Version 0.12.9) and the Illumina reads were mapped to the reference file using the recombinant-mapping algorithm, ViReMa (Viral-Recombination Mapper, Version 0.15). The recombinant sequences were compiled as a text file and analysed using ggpubr (v2.3) in R Studio.

## Results

### Modular RG system design for DWV

To compare the virulence and competitiveness of DWV types and their recombinants a set of cDNA clones were prepared. By exploiting the modular organisation of the DWV genome [21] we have previously constructed infectious cDNAs for several distinct genetic variants of DWV [25]. For convenience these are referred to as follows: VDD (DWV type A coding sequence, GenBank MT415949), VVD (a type B/A recombinant, GenBank MT415950) and VVV (DWV type B, GenBank MT415952). In addition we constructed a cDNA for a complete type A DWV, designated DDD, using a similar gene synthesis and module replacement strategy [25] to incorporate the DWV type A 5’-untranslated region (5’-UTR; DWV-A 1414, GenBank KU847397 used as a reference - Figure S1). VDD, VVV and VVD DWV variants were previously shown to be infectious and cause symptomatic disease in honey bees [25]. Infectivity of the DDD virus was verified by analysis of DWV accumulation in injected pupae and was indistinguishable from the VDD virus (Figure S2a). Derivatives of VDD, VVD and VVV, expressing the enhanced green fluorescent protein (EGFP) or mCherry, were generated as previously described [25] (Figure S1) and their replication verified following inoculation of pupae (for example, Figure S2b).

### Superinfection and coinfection studies

*Varroa* delivers DWV to developing honey bee pupae by direct injection when feeding. Pupae will already contain previously acquired DWV and the mite may contain one or more DWV variants. We investigated the consequences of coinfection and superinfection on accumulation of distinct DWV variants in honey bee pupae under laboratory conditions. Primary infection was achieved by feeding first instar larvae (0-1 day old) with a diet containing 10^7^ genome equivalents (GE) of either VDD or VVV DWV, followed by secondary inoculation by injection (10^3^ GE) with the reciprocal virus variant ten days later at the white-eyed pupal stage. The viral load in individual pupae was analysed by qPCR 24 h post-injection using DWV type-specific primers for the RNA-dependent RNA polymerase (RdRp) coding region. Pupae infected by larval feeding showed a markedly reduced accumulation of the injected DWV variant when compared to the same virus in pupae which were not fed DWV as larvae (Figure 1a).

**Figure 1.**
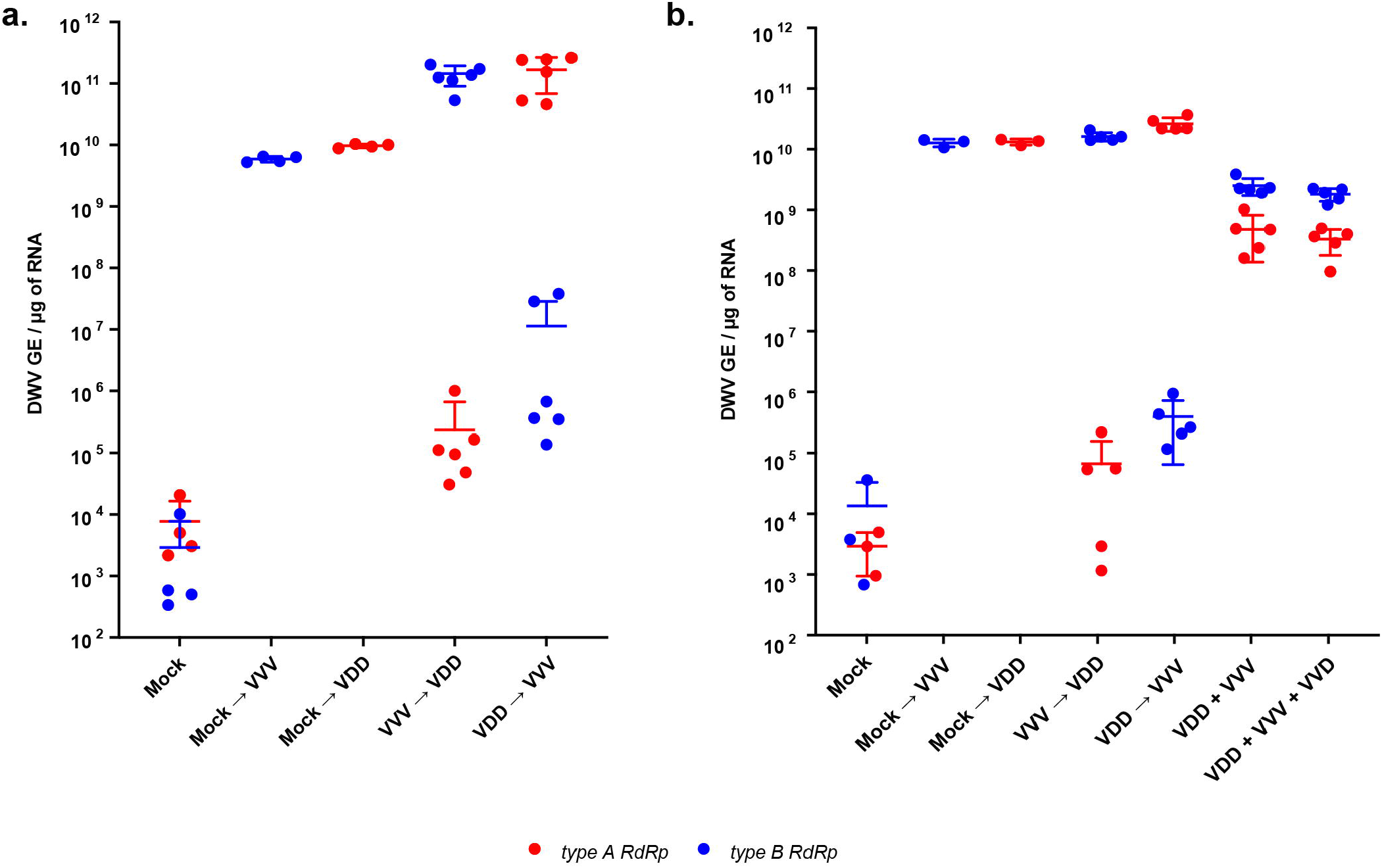
Coinfection and superinfection of honey bee brood with DWV. **a.** Superinfection of honey bee pupae preliminarily infected *per os* at larval stage and then via injection at pupal stage. Quantified viral titres of VVV and VDD DWV in pupae 24 h post-injection of the superinfecting virus are shown (second virus inoculated by injection after primary infection by feeding with the reciprocal DWV variant is indicated by “→”, *e.g.* VVV (fed) → VDD (injected)). **b.** DWV accumulation in honey bee pupae in coinfected (mixed virus population indicated by “+”) or superinfected samples (second injection 24 h after primary infection by injection is indicated by “→”). Primer pairs for type A or type B RdRp amplifying variant-specific fragments of virus polymerase encoding region were used to distinguish between the administered variants. Data points represent DWV levels in individual pupae with two points of different colour corresponding to different virus variants (red for type A RdRp and blue for type B RdRp respectively) in the same pupa (or in individual pupae for VVV or VDD only injected samples). Error bars show mean ±SD for each virus variant in each injection group, GE - genome equivalents. ANOVA: P<0.05 for type A accumulation in “Mock→VDD” vs “VVV→VDD” and for type B level in “Mock→VVV” vs “VDD→VVV” groups.

Reduced accumulation of a superinfecting virus was also observed when white-eyed pupae were initially injected with VDD or VVV 24 h prior to introduction of the reciprocal virus variant (first injection - 10^2^ GE, superinfection - 10^6^ GE, Figure 1b). In contrast, simultaneous infection with two or three (VDD, VVV and VVD) DWV variants (10^2^ GE in total virus injected corresponding to 0.5×10^2^ or 0.33×10^2^ GE of each variant for two- and three-component infections respectively) resulted in nearly equivalent virus loads, although the VDD variant accumulated to slightly lower (~0.5 log_10_) titres at 24 h post-injection.

### Dynamics of DWV accumulation in superinfection conditions

We extended these studies to determine whether the apparent competitive disadvantage for the second virus remained after an extended incubation period. Pupal injections were repeated as before and viral loads quantified 5 and 7 days after superinfection. A recombinant type B/A variant (VVD) was additionally included both as primary and superinfecting virus. In reciprocal infections using VDD and VVV both the initial and the superinfecting virus reached nearly equivalent levels within the incubation period (Figure 2). In contrast, in virus pairings with a greater sequence identity between the genomes the superinfecting virus exhibited a reduced accumulation even after prolonged incubation. In the “VDD→VVD”, “VVD→VDD” and “VVD→VVV” groups the superinfecting virus levels were ~2 log_10_ lower than the initial inoculum at 5-7 days post-injection. For the “VVV→VVD” pairing this was more marked, with the superinfecting virus still ~4 log_10_ lower after 7 days. In control pupae infected with VDD, VVD or VVV individually all three viruses reached high titres 7 days post-injection (Figure 2 and Figure S3). Additionally, virus accumulation was monitored after coinfection of equal amounts of each combination of VDD, VVD and VVV over time. In these studies, all coinfecting variants achieved similar titres 5 days post-inoculation (Figure S3).

**Figure 2.**
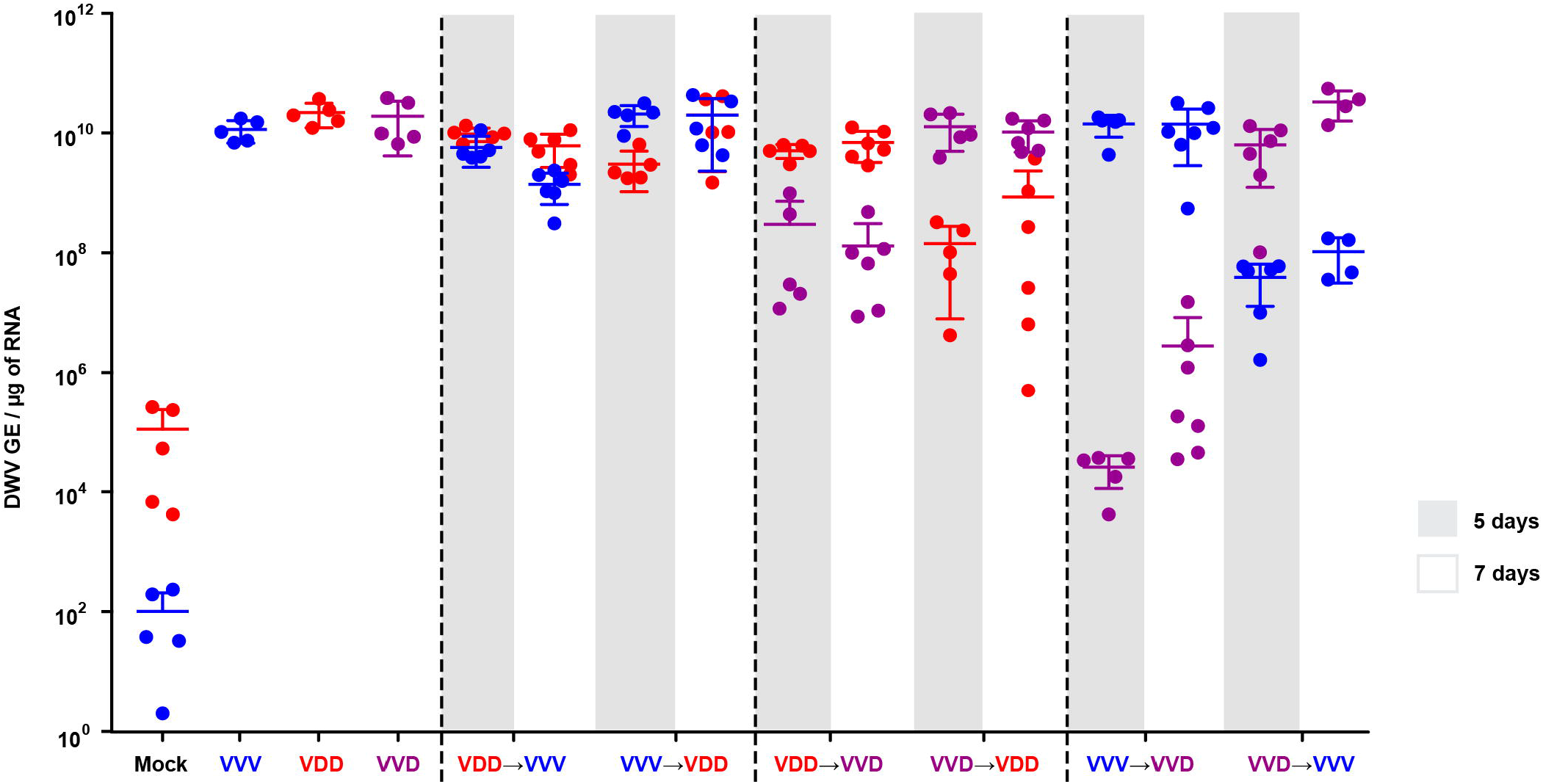
qPCR analysis of DWV accumulation in superinfection conditions. Honey bee pupae received a primary injection with one DWV variant (VVV, VDD or VVD) and a secondary injection (superinfection) with a different variant 24 h later. DWV accumulation was quantified 5 (grey shading) and 7 (no shading) days after the second injection. Primer sets specifically targeting the viral polymerase or a structural protein encoding region of DWV type A or type B were used to detect accumulation of each of the injected variants. Data points represent DWV levels in individual samples with two points of different colour corresponding to different virus variants (red for VDD, blue for VVV and purple for VVD respectively) in the same pupa (or in individual pupae for VVV, VDD or VVD only injected samples). Error bars show mean ±SD for each virus variant in each injection group, GE - genome equivalents. ANOVA: P<0.05 for “Mock→VDD” vs “VVD→VDD”, “Mock→VVD” vs “VDD→VVD”, “Mock→VVD” vs “VVV→VVD”, “Mock→VVV” vs “VVD→VVV” at 7 days time point.

We recently demonstrated that bumble bees are susceptible to DWV infection when directly injected at high doses [38]. We therefore investigated the influence of the host environment on the DWV superinfection by conducting similar experiments in bumble bee pupae. At 48 h post superinfection the levels of the second virus administered were lower than that of the primary virus inoculated but – with the exception of the “VVD→VVV” combination – had achieved similar levels by 6 days post-injection (Figure S4).

These results suggest that a superinfecting virus experiences an initial competitive disadvantage, but that this disadvantage is overcome after 5 to 7 days unless the viruses exhibit more extensive sequence identity. To investigate this further we studied superinfection with essentially identical viruses, using two VVD variants distinguishable solely by unique genetic tags – VVD_S_ and VVD_H_, tagged with a *SalI* or *HpaI* restriction site respectively (Figure S1) – which differ by just 4 nucleotides. Honey bee pupae injected with VVD_H_ were challenged 24 h later with VVD_S_ and analysed by end point PCR and restriction assay 1, 3 and 6 days after superinfection. No VVD_S_ was detectable in superinfected pupae at any time point analysed (Figure S5) suggesting a complete or near-complete block of the superinfecting genome amplification. Control injections of VVD_S_ into pupae, which did not receive VVD_H_ virus, allowed detection of *SalI*-tagged cDNA 24 h post-inoculation.

### Tissue localisation studies using reporter-encoding DWV

Total RNA levels analysis allows the quantification of DWV to be determined, but it obscures details of the relative distribution and tissue tropism of individual virus variants. Previously we developed an EGFP-encoding RG system for DWV [25] based upon the VDD genome and designated DWV_E_ (for convenience here renamed to VDD_E_). We used VDD_E_ to define whether the primary infection also affects the distribution of the superinfecting virus. Furthermore, we constructed a full length DWV type A genome, designated DDD (Figure S1), and similarly investigated superinfection of DDD infected pupae. Pupae that had received an initial injection of 10^2^ GE of DDD, VDD, VVD or VVV were inoculated 24 h later with 10^6^ GE of VDD_E_. Live pupae were analysed by confocal microscopy for the presence of the EGFP signal (Figure 3). Three regions of each pupa were visualized - the head, the developing wing and the abdomen - as we have previously demonstrated significant virus accumulation in these locations [25].

**Figure 3.**
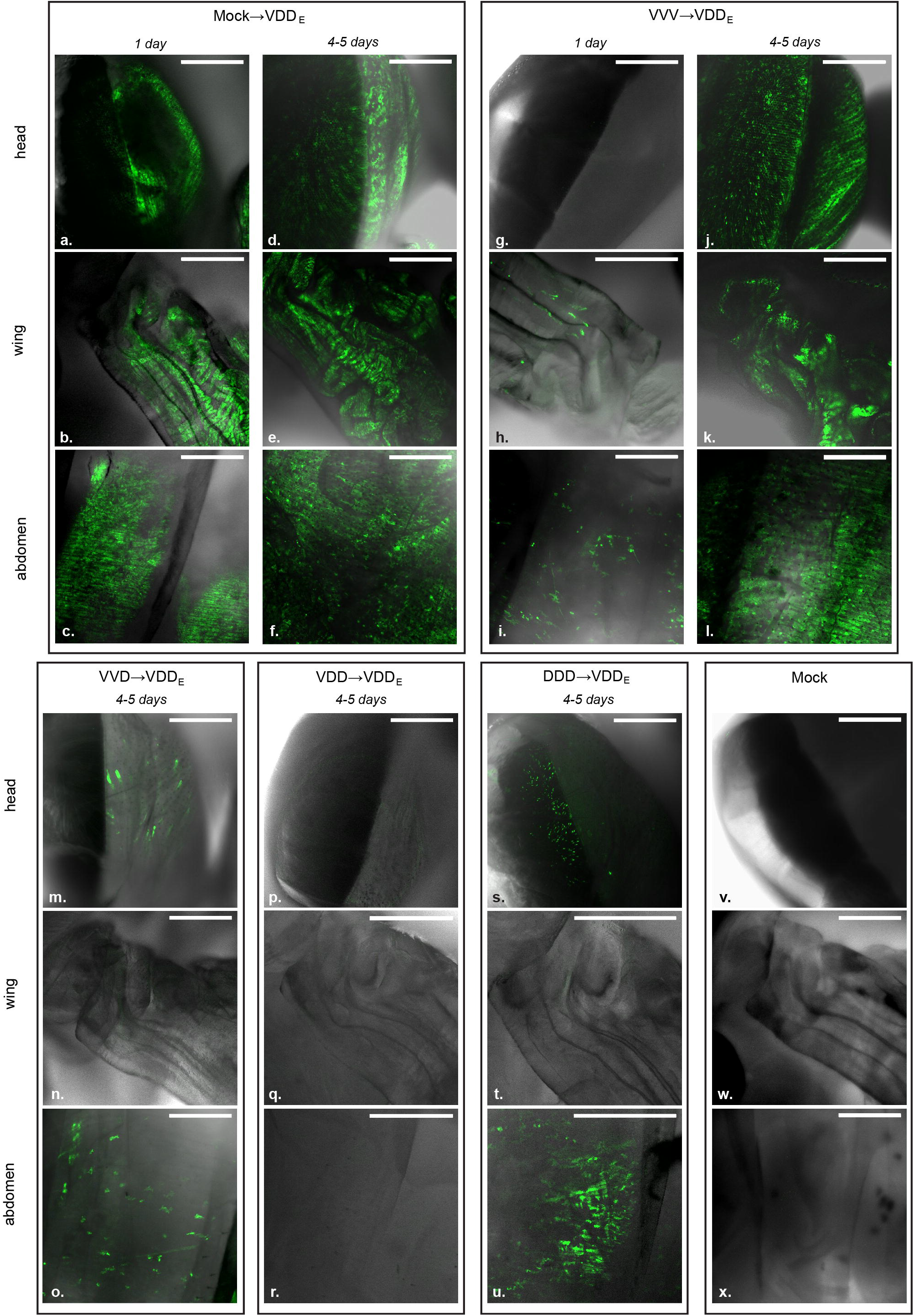
EGFP signal localisation in VDD_E_ injected honey bee pupae analysed by confocal microscopy. Combined white-field and fluorescent images are shown for convenience of interpretation. Pupae were analysed 1 and 4-5 days after the second injection. For VVD, VDD and DDD primary infection groups only samples incubated for 4-5 days are shown, as no EGFP was detected after 1 day. Scale bars correspond to 500 μm.

Injection of VDD_E_ in the absence of a primary infection (“Mock→VDD_E_” group) resulted in efficient expression of EGFP throughout the pupa 24 h post-inoculation (Figure 3a-c). In the case of superinfection, the EGFP signal could be seen 24 h later only in pupae where VVV was used as a primary infecting genotype (Figure 3h and i). In these pupae, the number of fluorescent foci was lower when compared to the “Mock→VDD_E_” group infected for the same 24 hperiod (Figure 3, panels a-c *vs*. g-i in). No EGFP signal was visible upon superinfection with VDD_E_ after 24 h in pupae first injected with VVD, VDD and DDD (data not shown). At 4-5 days post-inoculation with VDD_E_ there were also differences observed in the levels and distribution of the reporter protein. For example, no EGFP signal was found in the wings after primary inoculation with VVD or DDD (Figure 3, panels n and t *vs.* e and k). Visible EGFP expression was detected in the head and abdomen in the pupae from these injection groups after 4-5 days but the extent and number of fluorescent foci was reduced when compared to the “Mock→VDD_E_” and “VVV→VDD_E_” pupae (Figure 3, panels m, o, s and u *vs*. panels d, f, j and l). In contrast to the “Mock→VDD_E_” and ”VVV→VDD_E_” samples, only a fraction of pupae in ”VVD→VDD_E_” and ”DDD→VVD_E_” groups exhibited detectable EGFP signal in each of the body sites under analysis (Table S2). Finally, pupae initially injected with VDD did not show any detectable EGFP signal even 6 days after superinfection with VDD_E_ (Figure 3p-r, Figure S6), suggesting again that greater sequence identity restricts the activity of the superinfecting virus.

To confirm that the external analysis of the intact living pupae was representative, selected samples were dissected. Tissue samples, including parts of the digestive tract, wing rudiments, thoracal muscle tissue, brain, and cephalic glands were visualised by confocal microscope (Figure S6). This analysis recapitulated the pattern of fluorescence observed by previous visualisation of intact pupae. To complement the microscopy data we quantified DWV RNA in selected pupae by qPCR at 24 h and 5 days post superinfection (Figure S7) and found that there was a good agreement between the amount of genomic RNA and the level of detectable fluorescence.

### Localisation of DWV in coinfected and superinfected pupae using two-colour microscopy

In order to visualize the distribution of infection with different DWV variants we used EGFP- and mCherry-expressing viruses, VDD_E_, VVD_E_ and VVV_mC_ (with subscript E and mC indicating the EGFP or mCherry reporter respectively, Figure S1). For coinfection, pupae were injected with equimolar mixtures of VVD_E_ or VDD_E_ and VVV_mC_ and analysed under the confocal microscope 1 to 5 days post-inoculation. We could readily detect red and green fluorescent signals present in the same tissues of virus-injected pupae as previously described [25], including multiple tissues of the digestive tract, wings and head tissues. The reporter gene expression sites appeared as individual punctate foci of either red or green fluorescence, with only a few displaying dual fluorescence for both reporters (Figure 4a and Figure S8). The analysis of VDD_E_–infected pupae superinfected with VVV_mC_ and visualised by microscopy after a further 24 h revealed a similar distribution of the fluorescent signal as in coinfected samples (Figure 4b).

**Figure 4.**
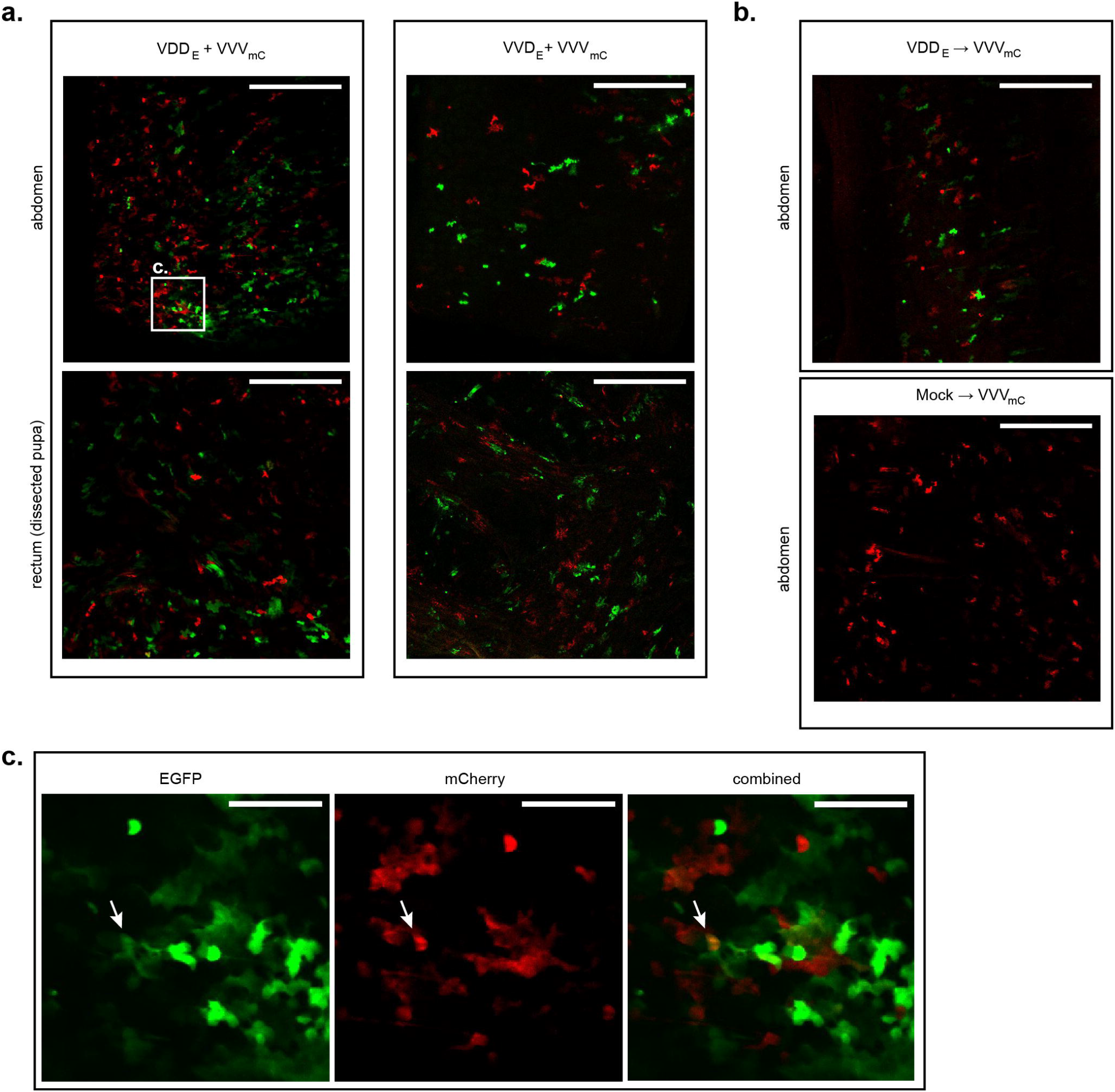
Confocal microscopy analysis of honey bee pupae coinfected or superinfected with DWV variants encoding EGFP and mCherry. **a.** Coinfection: “VDD_E_ + VVV_mC_” or “VVD_E_ + VVV_mC_” panels show red (mCherry) and green (EGFP) fluorescence signals detected in the abdomen of intact pupae (upper row) or in the dissected tissues of the digestive tract (rectum tissue shown as an example). Scale bars correspond to 500 μm. **b.** Superinfection: upper panel – abdomen of an intact pupa initially infected with VDD_E_ and superinfected 24 h later with VVV_mC_ analysed 24 h after the second injection; lower panel – abdomen of an intact pupa injected with VVV_mC_ only and analysed 24 h post-inoculation. Scale bars correspond to 500 μm c. Magnified image of highlighted region (from panel a) of pupa coinfected with VDD_E_ + VVV_mC_. Individual images for EGFP and mCherry signals, and a combined image for both fluorophores are shown. Arrows indicate Individual foci of infection exhibiting both EGFP and mCherry expression. Scale bars correspond to 100 μm.

### Recombination between VDD and VVV DWV

The interpretation of the superinfection studies is based upon sequence-specific quantification of particular regions of the virus genome by qPCR. This interpretation could be confounded by extensive levels of genetic recombination, a natural consequence of coinfection with related viruses [40]. Genetic recombination of RNA viruses requires that both parental genomes are present within an individual cell [41]. Since our microscopy analysis had detected only limited numbers of apparently dually infected foci during mixed infections we conducted further analysis to investigate the presence and identity of viral recombinants, and the influence of the order of virus acquisition on recombination, using next generation sequencing. Illumina paired-reads were generated from PCR amplicons of a 10 Kb fragment of the DWV genome targeting pupae initially infected with VDD and challenged with VVV, or vice versa (samples “VVV→VDD” and “VDD→VVV” at 5 or 7 days after superinfection). Recombination junctions were detected across the entirety of the DWV genome in all samples analysed with ‘hotspots’ of recombination denoted by an increased number of aligned reads identified at numerous points in the genome (Figure 5b, Table S3), including some previously reported [21]. The percentage of reads corresponding to recombination junctions varied in individual pupae from 1-2.2% of all mapped reads (Figure S9). In all cases approximately equal proportions of recombinants were detected with VVV or VDD as the 5’-acceptor partner (terminology assumes that recombination occurs during negative strand synthesis [40, 42]). In several instances we detected the same recombination junction with both VVV and VDD as the 5’-acceptor. Our analysis also revealed recombination sites in which the 5’ was only ever derived from one variant or the other (red and blue points in Figure 5b, Table S3). These results demonstrate that although superinfecting virus recombines readily with an established variant, the recombinant population remains a minor component of the total virus population, and is well below the level expected to confound our analysis of competition between extant and superinfecting viruses.

**Figure 5.**
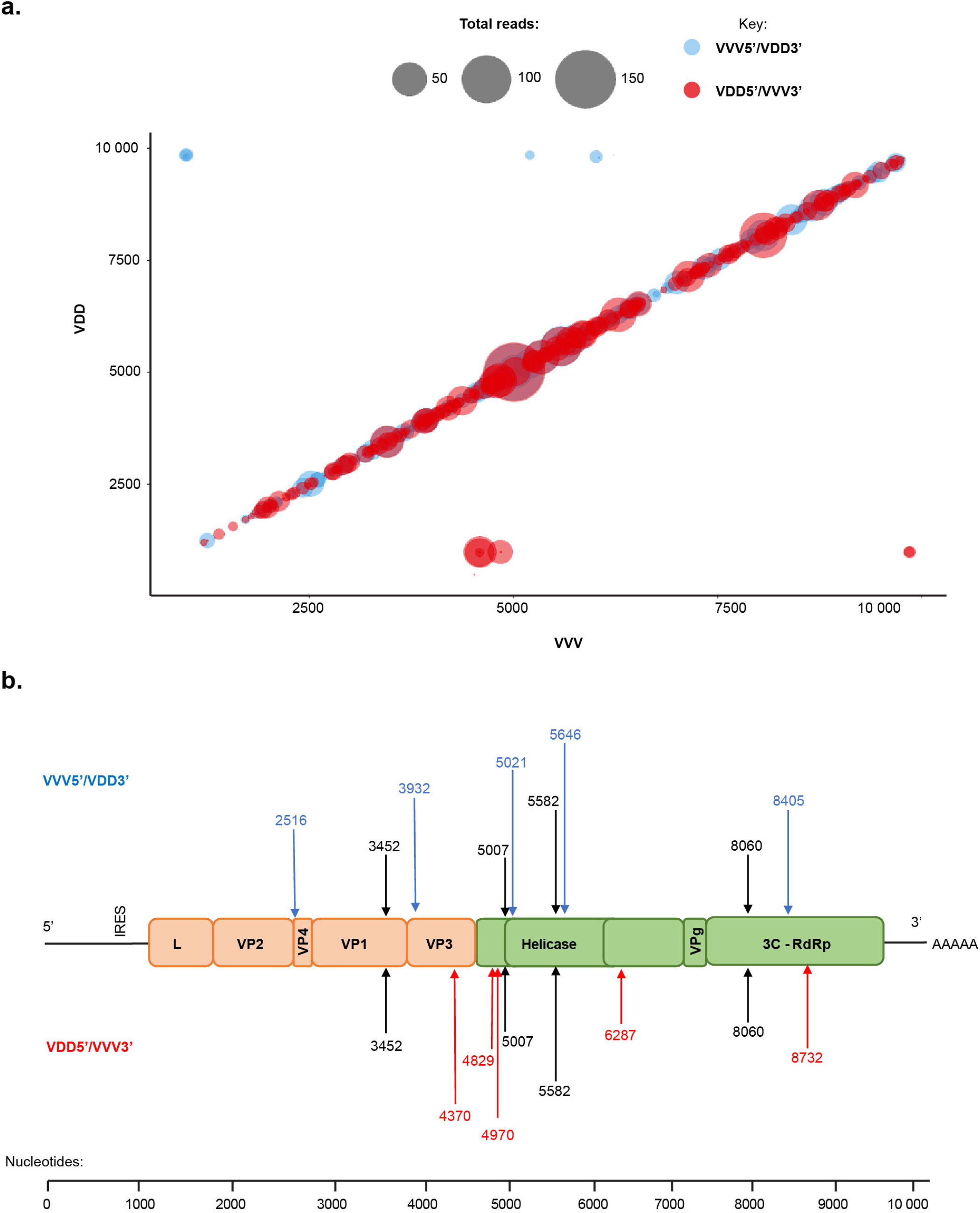
Genomic recombination events observed between VVV and VDD DWV variants in a superinfected honey bee pupa. **a.** Mapped recombination events in a honey bee pupa initially infected with VVV and superinfected with VDD DWV. The plot shows recombination events occurring along the full length of the DWV genome, with the VDD genome length shown on the Y-axis and VVV shown on the X-axis. Each bubble represents a unique recombination site and bubble size is determined by the number of mapped and aligned reads for this site obtained using ViReMa analysis. The colour of the bubble indicates the direction of recombination, with blue representing VVV as 5’-acceptor and VDD as 3’-donor and those in red with VDD as 5’-acceptor and VVV as 3’-donor. **b**. The most frequently observed recombination junctions in all pupal samples analysed, shown at the recombination junction between VVV and VDD genomes. The junctions shown in black occurred with similar frequency in both directions. Those shown in blue occurred predominantly with VVV as 5’-acceptor and VDD as 3’-donor and those in red with VDD as 5’-acceptor and VVV as 3’-donor sequences.

## Discussion

The global distribution and ubiquitous nature of DWV [14, 19], transmitted vertically and horizontally in honey bees [15, 43, 44], inevitably means that when vectored by *Varroa* it is introduced to the host as a superinfecting virus. As such, there is the potential for competition for cellular resources in coinfected tissues, or the possibility of a pre-existing infection retarding or inhibiting superinfection through molecular mechanisms including SIE or the immune responses induced by the initial virus. There are at least two distinct types of DWV circulating globally – type A and type B – with documented differences in their distribution [8, 18, 22] and, perhaps, pathogenesis [13, 19, 23, 25]. If the outcome of superinfection always favoured one virus type it would influence transmission of DWV variants potentially accounting for their geographic distribution and – if associated with differences in virulence – the impact on the honey bees.

SIE has been reported for DWV, with the suggestion that bees bearing a type B virus were protected from subsequent type A transmitted from infesting *Varroa* mites [10]. SIE is described for several human, animal and plant viruses [26–35], and may operate via a number of molecular mechanisms [26, 27, 30, 31, 45–48]. Precedents already exist in plants with milder forms of a virus providing protection against more virulent strains [49, 50] and the recent spread of DWV type B in the USA [8, 22] could be interpreted as an indirect consequence of SIE, with bees harbouring this virus less susceptible to infection by DWV type A. However, there are other potential differences between DWV types such as the ability of variants with type B capsid to replicate in *Varroa* [25, 51], which may enhance its spread over the non-propagative transmission reported for type A [52].

The availability of RG system allowed us to investigate the consequences of coinfection and superinfection with DWV type A and B in individual honey bees. We found that when coinfected DWV type A and B (VDD and VVV variants) demonstrate broadly similar levels of replication (Figure 1b). In contrast, in sequential infections, either of virus-fed larvae or injected pupae, superinfecting DWV variant showed delayed replication (Figure 1). This delay was dependent upon the genetic similarity of the primary and secondary viruses and appeared transient in certain pairings. In genetically divergent pairings (*e.g.* “VDD→VVV” and “VVV→VDD”) high levels of both viruses were reached after a prolonged incubation period. In contrast, where the extent of genetic identity between the primary and secondary virus was greater, the superinfecting virus failed to ‘catch up’, even after 7 days (Figure 2). This was most dramatically demonstrated using two genomes that differed by just 4 nucleotides (VVD_S_ and VVD_H_ variants), in which case the superinfecting virus remained undetectable after 6 days incubation (Figure S5). In addition, we found that delayed accumulation of the genetically similar superinfecting DWV variants is not specific to honey bee host and was also observed in bumble bees, a species susceptible to DWV infection when directly injected (Figure S4, [38]).

We extended our analysis in honey bee pupae using reporter gene-expressing viruses and demonstrated that replication, characterized by the expression of the fluorescent protein, was inversely related to the level of genetic identity between the primary and superinfecting viruses (Figure 4). For example, VDD_E_ replicated extensively, albeit somewhat delayed when compared with VDD_E_-only infected pupae, in pupae that had received VVV as the primary virus (Figure 4j-l), but was undetectable in pupae initially inoculated with VDD (Figure 4p-s). Notably, in each case where the superinfecting virus showed reduced replication after extended incubation, dominance in the replication showed no directionality according to virus type and was due solely to the order of addition. Based on this data it is likely that sequential infection with DWV type A and B will result in both viruses replicating to maximal levels before eclosion of either worker or drone brood pupae (which pupate for ~12 or ~14 days respectively). It remains to be determined whether the delay we demonstrate is sufficient to influence the colony-level virus population, or that carried and transmitted by *Varroa*.

Where cellular coinfection occurs viruses have the opportunity to genetically recombine. This is a widespread phenomenon in the single-stranded positive-sense RNA viruses [53, 54] and has previously been documented in DWV [6–9]. Our microscopy analysis of honey bee pupae infected with two reporter-expressing DWV variants predominantly demonstrated non-colocalised expression of the fluorescent signal. However, small numbers of dual-infection foci were detected, directly implying that the opportunity for recombination exists (Figure 4c and Figure S8). Using next generation sequencing we confirmed the formation of recombinants and characterised the recombination products by analysis of the viral RNA in pupae reciprocally superinfected with VDD and VVV. 1-2.2% of mapped reads spanned recombination junctions, with no evidence for any bias in their directionality (VDD/VVV or VVV/VDD; Figure S9). Although these junctions mapped throughout the DWV genome, the greatest number were concentrated in the region of the genome encoding the junction of the structural and non-structural proteins (Figure 5a). This observation matches that found for other picornaviruses and reflects the mix’n’match modular nature of the *Picornavirales* genome. In this, functional capsid-coding modules can, through recombination, be juxtaposed with non-structural coding modules from a different parental genome [55]. A small number of recombination junctions (~350 of 35750 unique junctions mapped) plotted as outliers from the diagonal of genome-length recombinants. Analysis of these sequences showed that the majority were out of frame deletions (Woodford, unpublished), and so incapable of replicating. Our studies using analogous approaches in other RNA viruses show that these types of aberrant products are not unusual and reflect the random nature of the molecular mechanism of recombination ([42, 56]).

The competition we demonstrate in sequential DWV infections appears to be guided by the amount of genetic identity between the viruses. This suggests it is most likely mediated via RNA interference (RNAi). In arthropods antiviral RNAi response acts via generation of short double stranded RNAs (siRNA) from virus RNA replication intermediates through cleavage by the enzyme Dicer. These are further used by the RNA induced silencing complex (RISC) to target the destruction of complementary sequences [55, 56]. Hence viral RNA genomes exhibiting greater identity are likely to generate higher numbers of cross-reactive siRNAs. Previous analysis of the RNAi population in DWV infected honey bees demonstrated that 75% of DWV-specific short RNA are 21/22 mers [21]. Although DWV type A and B exhibit ~85% genetic identity it is not contiguous (Figure S1), but is instead distributed in ~1350 short regions of 1-389 nucleotides. Of these, less than 4% by number are of 21 nucleotides or greater in length, and therefore capable of generating perfectly complementary siRNAs. Recalculation of the identity between genomes having excluded sequences under 21 nt in extent demonstrates that there is only 34% genetic identity between DDD and VVV (Table S4). Comparing the figures from this analysis and the quantification of DWV accumulation in superinfected pupae suggests a clear relationship between the extent of the competition observed and the genetic identity of contiguous sequences. It is already known that exogenous RNAi can control DWV and other RNA viruses [57–59], and in our preliminary studies we have shown that RNAi-mediated suppression of Dicer leads to both increased pathogenesis and viral loads in DWV-infected bees (Figure S10). Further research will be required to determine the role of RNAi in competition between superinfecting DWV variants and its potential exploitation in studies to develop cross-reactive vaccines against DWV [36]. These future studies will need to take account of the disrupted complementarity between the genomes (Table S4), the uneven distribution of mapped RNAi’s to the genome [21] and both the variation acceptable within the RNAi seed sequence and the RNA structure of the target.

## Acknowledgements

We express our gratitude to Dr Marcus Bischoff and Gill McVee (University of St Andrews) for helping with microscopic imaging, Ashley Pearson (University of St Andrews) for assistance in molecular biology assays.

This work was supported by grant funding from BBSRC BB/M00337X/2 and BB/I000828/1.

