## Supplementary materials for "First come, first served: Superinfection exclusion in Deformed wing virus is dependent upon sequence identity and not the order of virus acquisition"

* corresponding author

### Supplementary materials

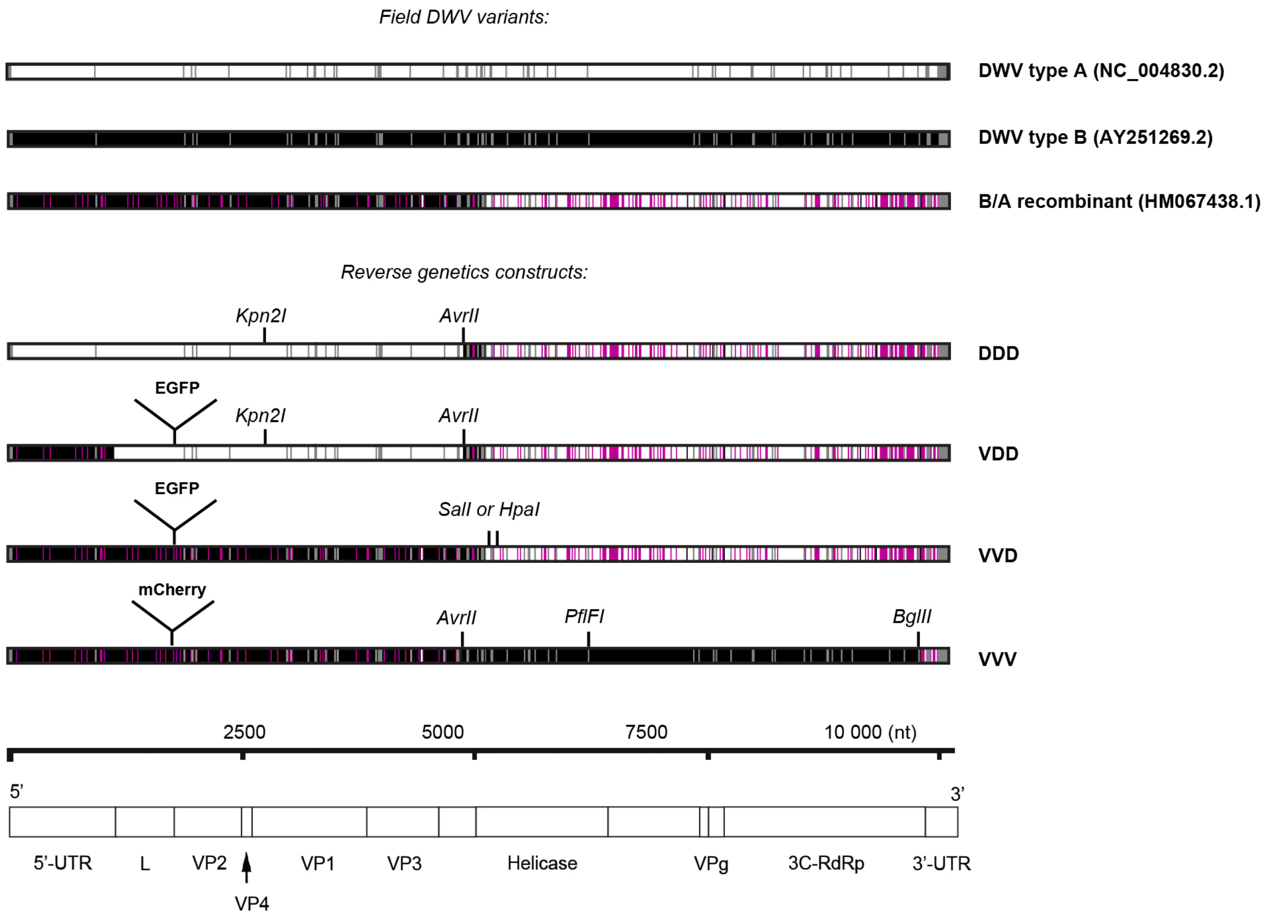


**Figure S1**. **Modular reverse genetics system design for DWV.** Diagram showing homologous parts between genomic RNA sequences of field DWV type A and B variants and the recombinant B/A clone used as a backbone for RG construction. DWV A and DWV B specific sequences are shown in white and black respectively, sequences unique to VDV-1-DWV-No-9 (GenBank HM067438.1) recombinant are shown in purple, identical regions between all three variants are shown in grey; positions of the new restriction sites introduced into each RG construct as genetic markers and location of reporter gene inserts are marked above the construct sequences; DWV genomic RNA organization is shown at the bottom: L - leader protein, VP1-VP4 - capsid proteins, VPg - virus protein genome-linked, 3C-RdRp - 3C protease and RNA-dependent RNA polymerase complex, 5' and 3'-UTR - flanking untranslated regions. VDD, VVD, VVV and DDD indicate constructed genomes based upon the modular structure of the DWV genome. Where fluorescent reporter-expressing derivatives were made (see [25] for indicative construction details) the name is suffixed with a subscripted E for EGFP-expressing or mC for mCherry expressing genomes, hence $VVD_E$ or $VVV_{mC}$. VVD variant cDNA was constructed in two versions – either containing *SalI* or *HpaI* restriction site as a genetic ‘tag’; these were further designated as $VVD_S$ and $VVD_H$ respectively.

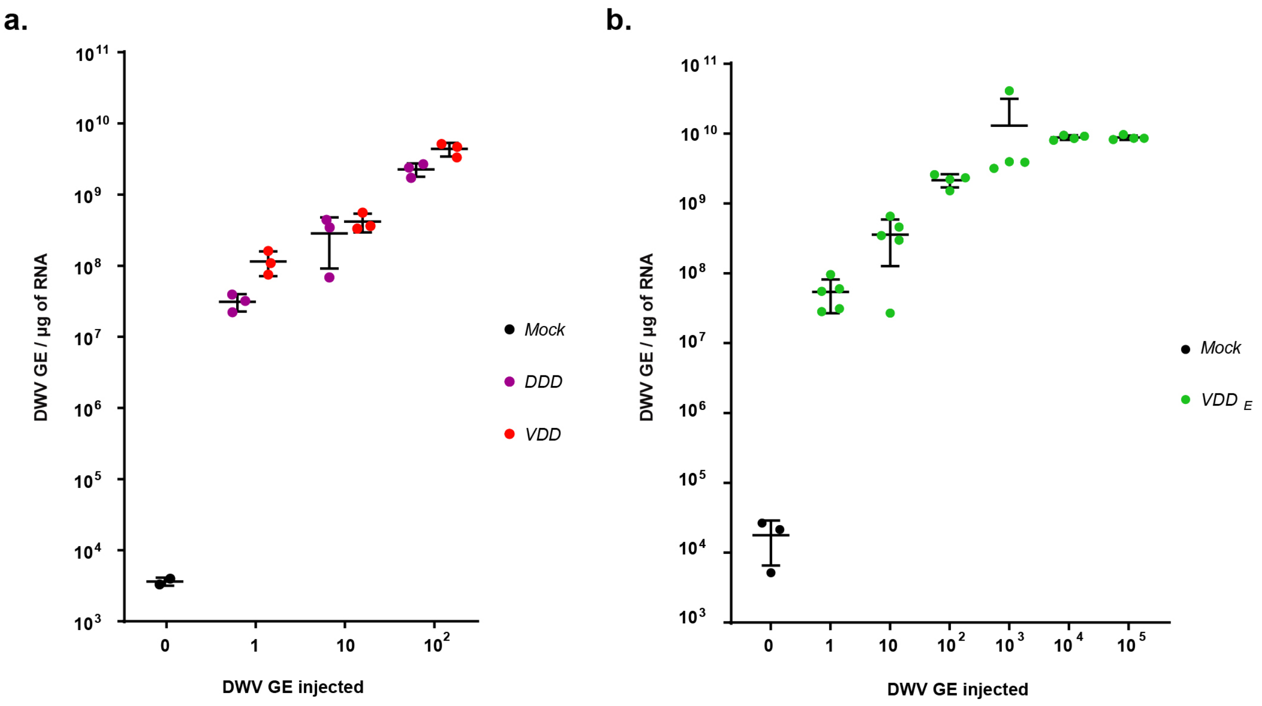


**Figure S2. qPCR analysis of DWV accumulation in injected honey bee pupae.** Pupae were injected at white-eyed stage and analysed 24 h post-inoculation, dots correspond to the virus level in individual samples, error bars show mean ±SD, GE - genome equivalents. **a.** Level of DWV in pupae inoculated with DDD or VDD virus. **b.** Level of DWV in pupae injected with $VDD_E$ virus.

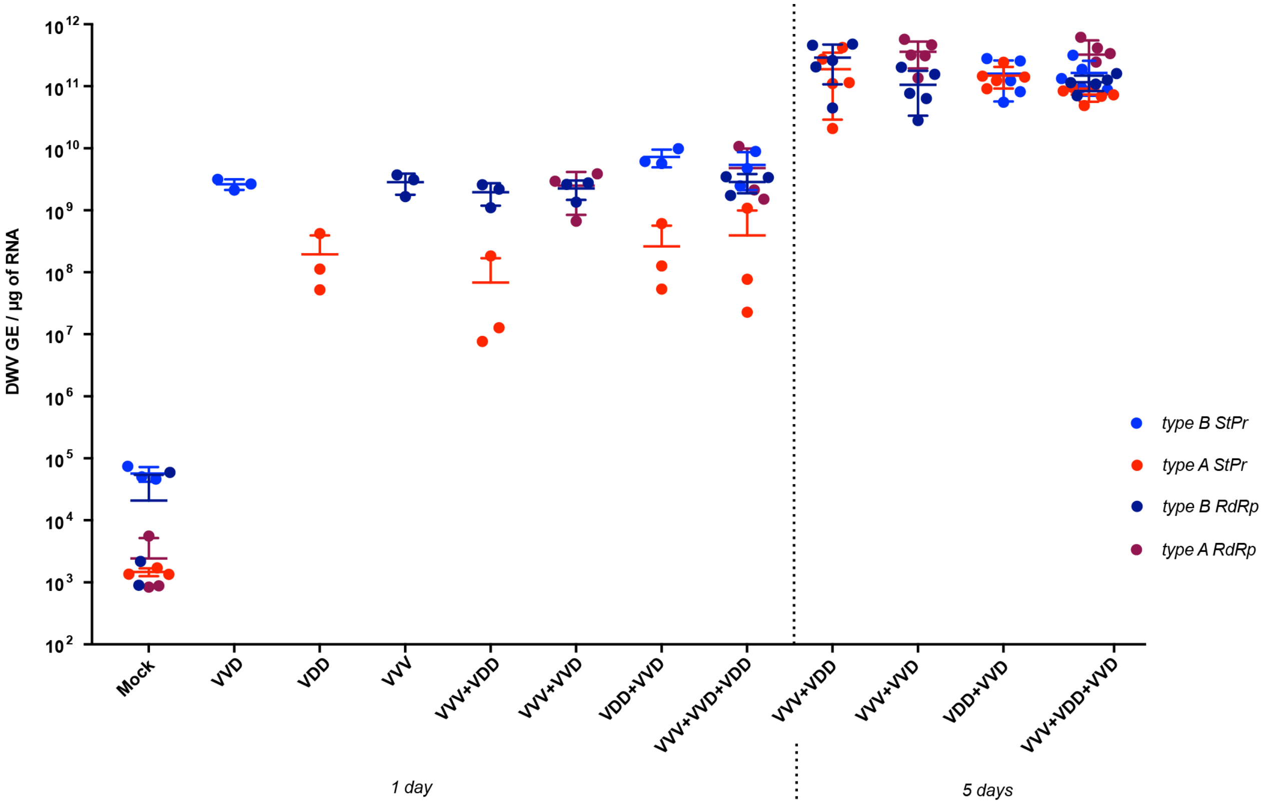


**Figure S3**. **Time course analysis of accumulation of DWV variants in honey bee pupae.** Mock (no injection) and virus-injected pupae were analysed 1 day and 5 days post-injection. Total amount of DWV administred per pupa was $10^2$ GE. Levels of specific genomic RNA were quantified individually in each sample using primer sets targeted to polymerase encoding region (RdRp primers) or structural proteins encoding region (StPr primers) of DWV type A and B. A StPr and B RdRp primers are specific to VDD and VVV DWV variants respectively, A RdRp primers target DWV type A cDNA region identical between VVD and VDD virus clones, B StPr primers target cDNA region identical between VVV and VVD variants. Each data point corresponds to an individual sample, in the case of mixed infection two or four (for three-component infection) data points indicating levels of each target are shown for each sample in the group. Eror bars show the mean level of each sequence variant quantified in the injected group ±SD, GE - genome equivalents.

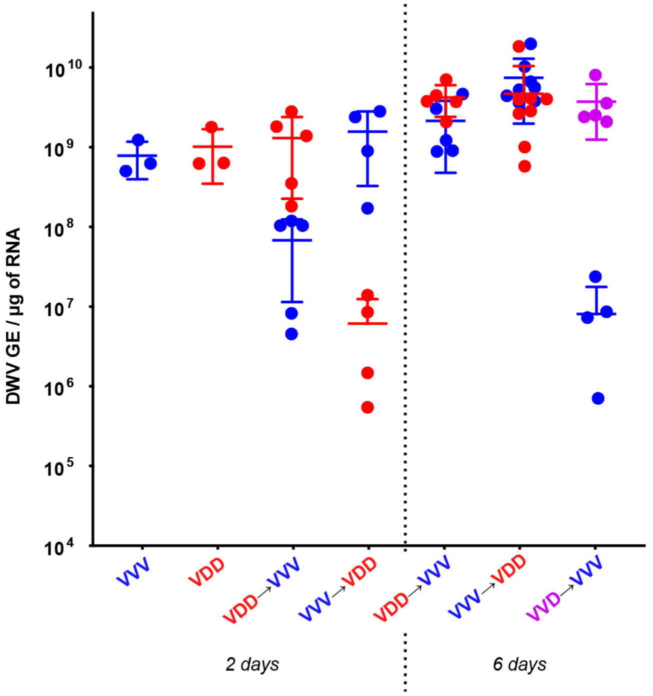


**Figure S4. Accumulation of DWV variants in superinfected bumble bee pupae.** Bumble bee pupae were inoculated with one variant of DWV (VVV, VDD or VVD; $10^3$ GE per pupa), then left for 2 days before being inoculated with a second variant ($10^6$ GE per pupa). Individual pupae were analysed after 2 or 6 days of incubation after superinfection by qPCR with type-specific primer pairs. Data points represent DWV levels in individual samples with two points of different colour corresponding to different virus variants (red for VDD, blue for VVV and purple for VVD respectively) in the same pupa (or in individual pupae for VVV, VDD or VVD only injected samples). Error bars show mean ±SD for each virus variant in each injection group, GE - genome equivalents.

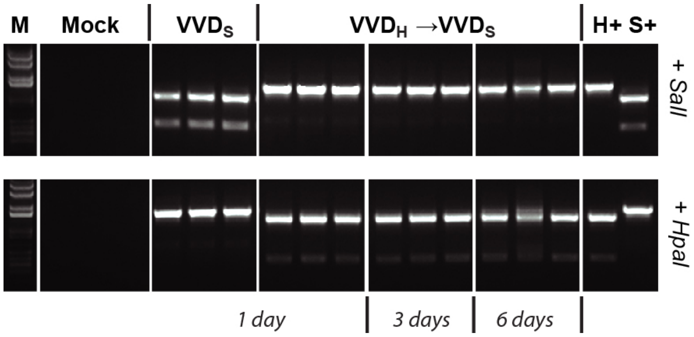


**Figure S5**. **Detection of DWV RNA with different restriction site tags by RT-PCR**. 1% agarose gel of restriction digest products shown. PCR samples in the upper panel were digested with *SalI*, those in the lower panel with *HpaI*. M - molecular size DNA marker, Mock - non-injected pupae samples, "$VVD_S$" - pupae, injected only with VVD virus tagged with *SalI* site (analyzed 1 day after infection); "$VVD_H \rightarrow VVD_S$" - pupae injected with $VVD_H$ (*HpaI* tagged VVD DWV) and superinfected with $VVD_S$ (*SalI* tagged VVD DWV) 1 day after the first injection and analyzed at 1, 3 and 6 days after superinfection. S+ and H+ are positive PCR controls for products containing *SalI* and *HpaI* sites respectively.

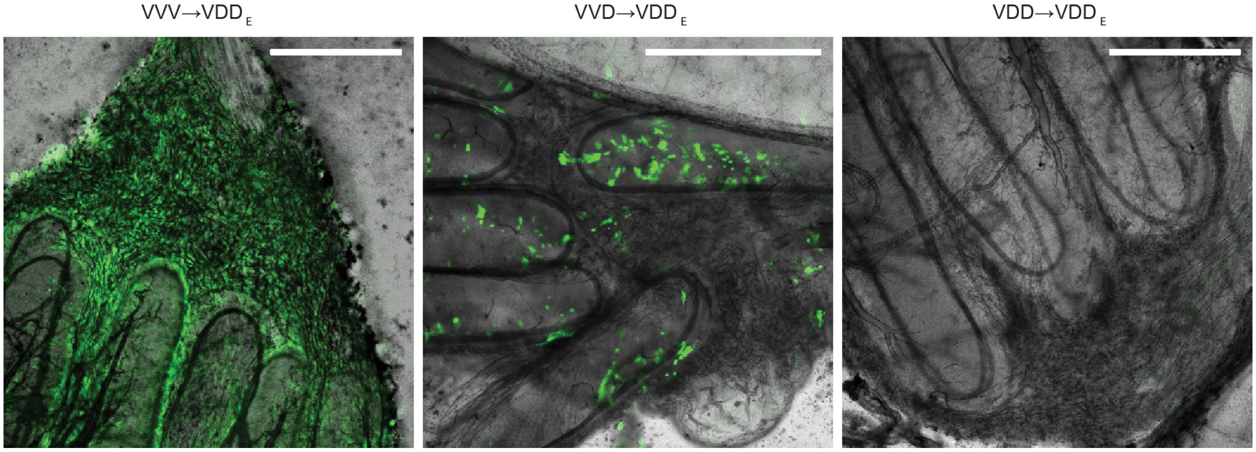


**Figure S6. Confocal microscopy analysis of dissected honey bee pupae injected with different DWV variants and superinfected with VDD$_E$.** Combined white-field and fluorescent images for rectum tissue of infected pupae are shown as examples. Pupae were analyzed 6 days after the second injection. Scale bars correspond to 500 μm.

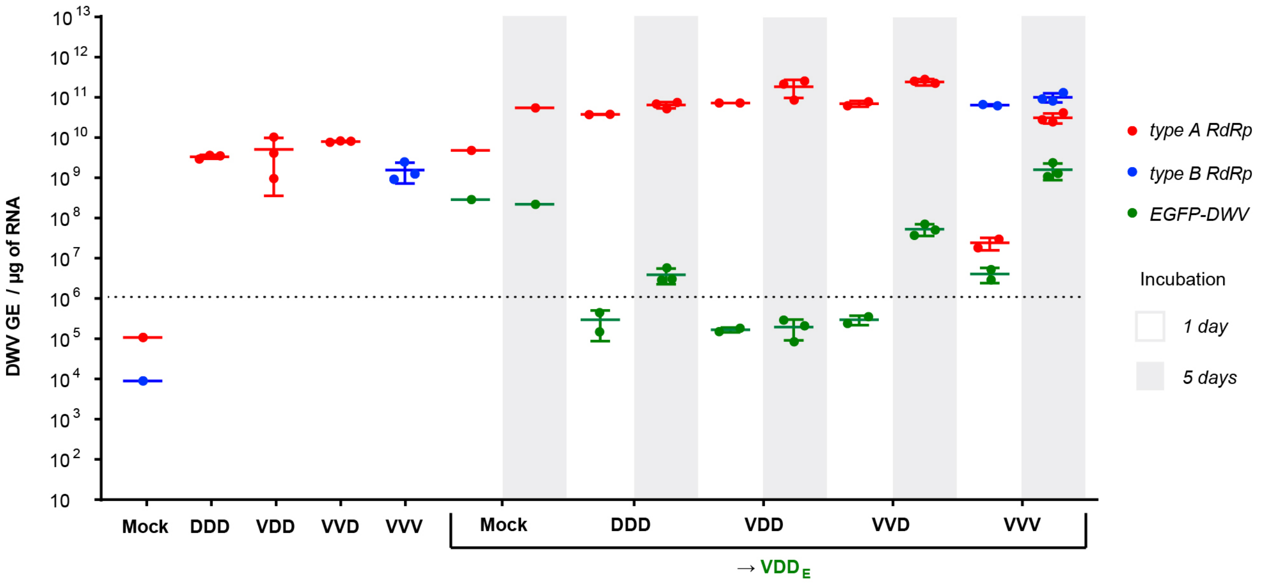


**Figure S7. Time-course analysis of DWV variants accumulation in honey bee pupae superinfected with VDD$_E$.** Pupae were analyzed 1 and 5 days after the last injection. qPCR analysis of the virus load was performed using primer sets targeting specific sequences encoding RNA polymerase of DWV type A (red dots) and type B (blue dots). Specific EGFP-encoding DWV RNA was detected by a primer pair targeting the junction between the EGFP insert and DWV genomic sequence (green dots). Each data point corresponds to an individual sample, in the case of mixed infection two or three data points indicating levels of each target sequence are shown for each sample in the group. Lines show the mean level of each sequence in the injected group ±SD, GE - genome equivalents. Individual pupae with VDD$_E$ RNA levels above the threshold indicated with the dotted line had visible EGFP signal when analysed by microscopy.

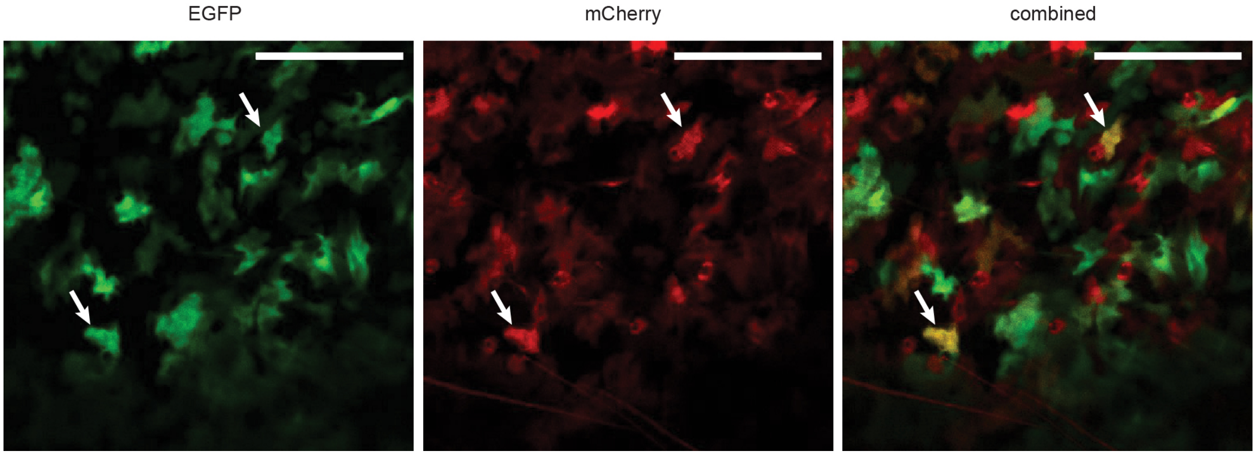


**Figure S8. Confocal microscopy analysis of a dissected honey bee pupa coinfected with DWV variants encoding EGFP ($VDD_E$) and mCherry ($VVV_{mC}$).** EGFP, mCherry and a combined image for both fluorophores obtained from a section of a digestive tract tissue are shown. Pupae were analyzed 3 days after injection. Individual foci expressing both fluorophores are indicated by arrows. Scale bars correspond to 150 μm.

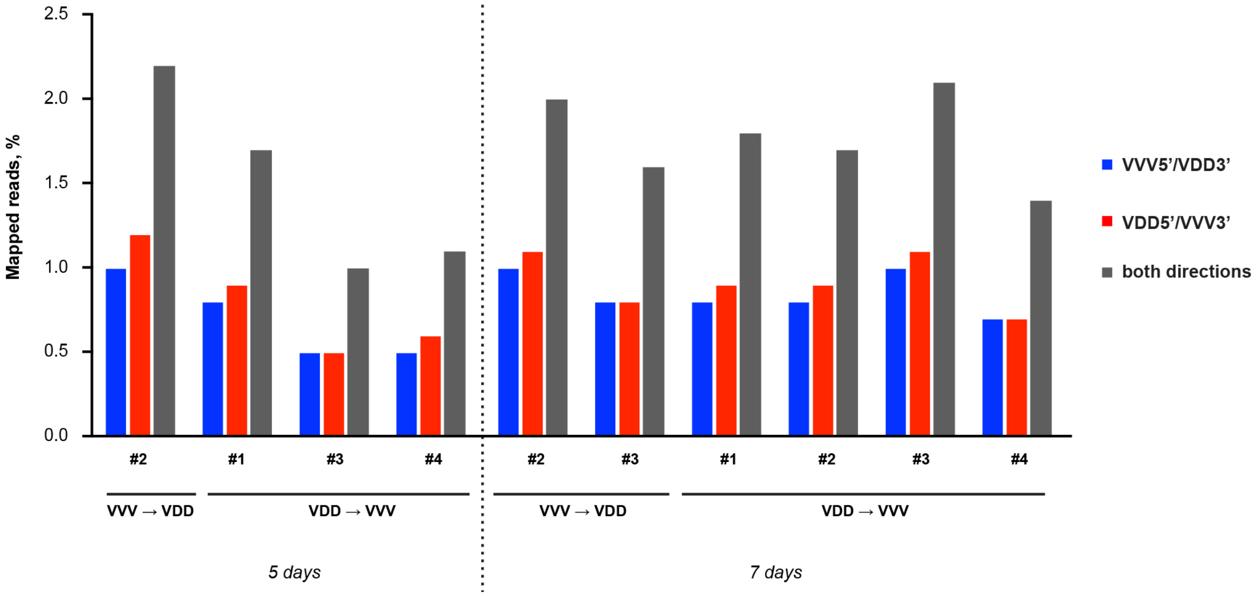


**Figure S9. Intertypic recombination reads in each of the honey bee pupae samples analysed using ViReMa and shown as a percentage of all mapped Illumina reads.** Blue and red bars represent amounts of recombinants with VVV5'/VDD3' and VDD5'/VVV3' directionality respectively, grey bars show the total amount of recombinants of both directionalities in each sample. Each cluster of bars corresponds to a single pupa sample with a sample number shown below the axis. "VVV→VDD" and "VDD→VVV" indicate samples from the corresponding superinfection injection groups and order of inoculation. 5 and 7 days refer to the incubation period of injected pupae after the superinfection injection.

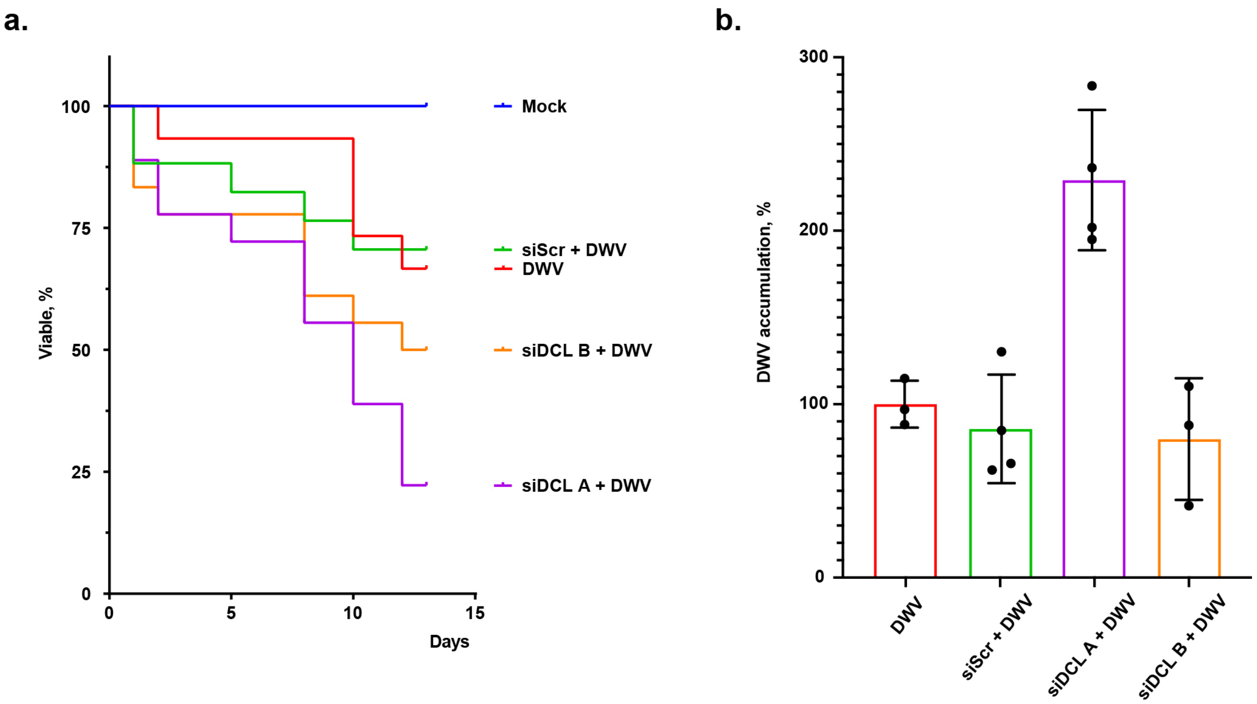


**Figure S10**: **Anti-Dicer siRNA treatment of honey bee larvae**. siRNA treatment was delivered in 5 µl of diet 3 times per day with 0.3-0.4 µg of siRNA supplied to each larva with each feeding and continued for 48 h after DWV inoculation (introduced together with 4th siRNA dose) or until pupation. “siDCL A” and “siDCL B” - larvae fed with specific siRNA targeting *Apis mellifera Dicer-like* mRNA (two different siRNA sequences were used), “siScr” - honey bee larvae fed with siRNA, which does not target any honey bee transcripts (control treatment), “DWV” - larvae fed with virus only (no siRNA treatment), “Mock” - non-treated control group; n=16 in each treatment group. **a.** Survival time course of honey bee larvae fed with siRNA and infected with VVD DWV. First instar honey bee larvae were treated with siRNA introduced with feeding from day one to day five of the experiment. On the second day larvae were orally infected with $10^6$ GE of VVD DWV. **b.** qPCR analysis of DWV accumulation in honey bee larvae fed with siRNA and orally infected with $10^5$ GE of VVD DWV. Larvae were analyzed 48 h after DWV feeding (72 h after the start of the siRNA treatment). Virus level in the DWV only group was set as 100%, each dot corresponds to DWV level in an individual larva sample, error barss representing mean ±SD.

**Table S1. PCR primers and oligonucleotides used in this study.**

| Name | Sequence (5'-3') | Target | cDNA binding site | Product size | Reference |
|---|---|---|---|---|---|
| **DWV_RTPCR_F** | ATTGATCATTGTATGTTTACCTTCCCTTG | DWV | 4021-4049 | 1606 bp | [25] |
| **DWV_RTPCR_R** | GCACGTAAGAGCTCGCTGCATA | DWV | 5606-5627 | | [25] |
| **DWV_qPCR_F** | ATATCACTTGGCGACGCAAC | DWV | 4950-4969 | 176 bp | [25] |
| **DWV_qPCR_R** | CCAATCTTTAAATTGTTTCGGTTTTTGAGC | DWV | 5098-5126 | | [25] |
| **DWV-A (RdRp) F** | TGTCTTCATTAAAGCCACCTGGAA | DWV type A | 8654-8677 | 140 bp | [13] |
| **DWV-A (RdRp) R** | TTTCCTCATTAACTGTGTCGTTGAT | DWV type A | 8769-8793 | | [13] |
| **VDV (RdRp) F** | TATCTTCATTAAAACCGCCAGGCT | DWV type B | 8654-8677 | 140 bp | [13] |
| **VDV (RdRp) R** | CTTCCTCATTAACTGAGTTGTTGTC | DWV type B | 8769-8793 | | [13] |
| **DWV type A StPr F** | GTCAAGAAGCAGGCGAATGTA | DWV type A | 1502-1522 | 129 bp | *This study* |
| **DWV type A StPr R** | GCATAGGGGATCTAGAACACATAG | DWV type A | 1608-1630 | | *This study* |
| **DWV type B StPr F** | GAAAAGACGCGGGTGAGTTCG | DWV type B | 1502-1522 | 130 bp | *This study* |
| **DWV type B StPr R** | AACATTGGCGATCGATTACAAACG | DWV type B | 1608-1631 | | *This study* |
| **Apis mellifera actin F** | AGGAATGGAAGCTTGCGGTA | *Apis mellifera actin* | 919–938 | 181 bp | [9] |
| **Apis mellifera actin R** | AATTTTCATGGTGGATGGTGC | *Apis mellifera actin* | 1099–1079 | | [9] |
| **EGFP_qPCR_F** | GAGAAGCGCGATCACATGGT | EGFP | 639-659 | 189 bp | [25] |
| **VDD_VP2qPCR_RP** | AGATGTACTAGGATCTCGCTGAGTT | DWV | 1860-1884 | | [25] |
| **mCherry_qPCR_F** | CAAGTTGGACATCACCTCCCAC | mCherry | 606-627 | 163 bp | [25] |
| **VVV_VP2qPCR_RP** | TAATTCAACTTCACCTTCGCCATCTG | DWV | 1808-1833 | | [25] |
| **DWV FG FP4** | GCGAATTACGGTGCAACTAAC | DWV | 19-39 | 10 068 bp | [38] |
| **DWV FG RP1** | TACGCGAGTAACACCTAAC | DWV | 10 076-10 094 | | [38] |
| **siScr1_FW** | GGATCCTAATACGACTCACTATAGTCTAGG ACCTCTGATATC | - | | | [60] |
| **siScr1_RV** | AAGATATCAGAGGTCCTAGACTATAGTGAG TCGTATTAGGATCC | - | | | [60] |
| **siScr2_FW** | GGATCCTAATACGACTCACTATAGATATCA GAGGTCCTAGAC | - | | | [60] |
| **siScr2_RV** | AAGTCTAGGACCTCTGATATCTATAGTGAG TCGTATTAGGATCC | - | | | [60] |
| **siDCL A top aintisense** | GGATCCTAATACGACTCACTATAGACTATG TACATAACGTGC | *Apis mellifera Dicer-like* | | | *This study* |
| **siDCL A bottom aintisense** | AAGCACGTTATGTACATAGTCTATAGTGAG TCGTATTAGGATCC | *Apis mellifera Dicer-like* | | | *This study* |
| **siDCL A top sense** | GGATCCTAATACGACTCACTATAGCACGTT ATGTACATAGTC | *Apis mellifera Dicer-like* | | | *This study* |
| **siDCL A bottom sense** | AAGACTATGTACATAACGTGCTATAGTGAG TCGTATTAGGATCC | *Apis mellifera Dicer-like* | | | *This study* |
| **siDCL B top aintisense** | GGATCCTAATACGACTCACTATAGATATAC ACCAATCAATGC | *Apis mellifera Dicer-like* | | | *This study* |
| **siDCL B bottom aintisense** | AAGCATTGATTGGTGTATATCTATAGTGAG TCGTATTAGGATCC | *Apis mellifera Dicer-like* | | | *This study* |
| **siDCL B top sense** | GGATCCTAATACGACTCACTATAGCATTGA TTGGTGTATATC | *Apis mellifera Dicer-like* | | | *This study* |
| **siDCL B bottom sense** | AAGATATACACCAATCAATGCTATAGTGAG TCGTATTAGGATCC | *Apis mellifera Dicer-like* | | | *This study* |

**Table S2. Percentage of samples with visible EGFP expression in each group of pupae superinfected with VDDE and analysed intact by confocal microscopy.**

| Injection group | Head | | Wing | | Abdomen | |
|---|---|---|---|---|---|---|
| | 1 day | 4-5 days | 1 day | 4-5 days | 1 day | 4-5 days |
| **Mock→$VDD_E$** | **100%** | **100%** | **100%** | **100%** | **100%** | **100%** |
| **VVV→$VDD_E$** | 0% | **100%** | **40%** | **100%** | **100%** | **100%** |
| **VVD→$VDD_E$** | 0% | **25%** | 0% | 0% | 0% | **88%** |
| **VDD→$VDD_E$** | 0% | 0% | 0% | 0% | 0% | 0% |
| **DDD→$VDD_E$** | 0% | **25%** | 0% | 0% | 0% | **38%** |

**Table S3. Positions of the most frequently observed recombination junctions in VVV and VDD DWV superinfected pupae when recombining in both directions, from VVV5'/VDD3' predominantly or VDD5'/VVV3' only.**

| Recombinant formed | Recombination point | Total mapped reads |
|---|---|---|
| | 5' nucleotide | |
| Both directions | 3452 | 600 |
| | 5007 | 1930 |
| | 5582 | 1179 |
| | 8060 | 789 |
| Predominantly VVV5'/VDD3' | 2516 | 246 |
| | 3932 | 257 |
| | 5021 | 326 |
| | 5646 | 266 |
| | 8405 | 242 |
| Only VDD5'/VVV3' | 4371 | 271 |
| | 4830 | 365 |
| | 4971 | 273 |
| | 6288 | 362 |
| | 8733 | 221 |

**Table S4. Genetic identity of RG DWV genomes.** DWV VVV, VVD, VDD and DDD genomes were aligned with ClustalX and the genome length genetic identity (percentage) calculated, as shown in the white cells in the upper right half of the table. Aligned DWV genomes were subsequently analysed for the percentage identity of sequences in excess of 20 nucleotides, as shown in the black shaded cells of the table. For example, VVV and DDD share 85% genetic identity, of which only 34% is located in contiguous sequences of more than 20 nucleotides.

| Identity | VVV | VVD | VDD | DDD |
|---|---|---|---|---|
| VVV | 100 | 93 | 87 | 85 |
| VVD | 68 | 100 | 94 | 92 |
| VDD | 41 | 73 | 100 | 98 |
| DDD | 34 | 67 | 93 | 100 |

**Text S1. Full cDNA sequences of DDD and $VVV_{mC}$ RG clones.** These constructs were obtained by modification of VDD (GenBank MT415949) and VVV (GenBank MT415952) cDNAs respectively. Gene-synthesis insert homologous to 5'-UTR sequence of DWV A (reference sequence - DWV-A 1414, GenBank KU847397) is highlighted in grey, mCherry encoding sequence is shown in red with flanking inserts encoding protease cleavage site duplication shown in bold font.

>DDD

tttaaaattcgctatgggaggcgatttatgccttccatagcgaattacggtgcaactaacaattttagata
gtagccataaacagacattatagtagctcactacgtattgatcattttataatgacttgcgtagtatgaagcgcat
gcttgtagttgtaactatgttactttgcaagttggagcttactattttggattatgaatatgtgcacttagtgtctg
tatttatagtcgtttgtggttcaaggttttgtgttagtagtacacttatgtatgaatgtacctttagtatgaatgtt
atagaatgacaatatcgaaggaaaaatctttataaaatacaaaaatattgtttttattatttcgatatggtgtttta
tagagtagattgccatgtgaccgctcatagaagtccattatggtttatcaatcgaagttgaatgtatttataagaat
attatacttaattagtaatattagtagtccgtaactattatcatccttttacagtttgatgtgataatagaccactg
cagtatcgagtagagtttcgaatgcgtagtgcaatagtacaatcactgtcaccgaccatctatcgtaatgatagatc
tgtcggaaaccattatttatgaagtgactagcaatcatggattaaattagatggtattctagtttagaggtgattcg
gcgctgcggtgcgactgaaacttctaaattagcatgtcagattgtattatgaatgcgttagtagtaatttctgcgat
agagctgggacccctcagtctctcaggtattgtatgaggcgaaagtgtgaaagttttgtatgtatttttttatatgt
acgactgtatcgggaattcctttagcaagaatccttttaatacagtataatctgtgctacggtacgttatgttcgca
gggcacccgttaatgtctcatagcccagacgatggcggatggaaagacatcatattttattttaatgctgtctttat
tgctgatttattttgctgtttttatttgctattttatatttgctaattttcattattgcgaaatatattatattgct
atttttattatatacgctagattcaattttattctttctatattttcaatttaattttgatttcgaaggtaaatata
tataattgattattaaaaatggcctttagttgtggaactctttcttactctgccgtcgcccaagctccgtctgtcgc
ctatgcacctcgtacatgggaagttgatgaagctaggcggcgccgagtcattaaacgtttggcgctggagcaagaac
gtattcgtaacgttcttgacgttgccgtctatgaccaggcgacatgggaacaggaggacgcgcgcgataatgagttc
ctaacggaacaattaaacaatttatatactatttattcgatcgctgaacgttgtacgcgtcggcctatcaaagagta
ctctcctatatcagtttcgaataggtttgctccactggaatccctcaaggtcgaggtcggtcaagaagcaggcgaat
gtatatttaagaaacctaaatatacgcgcgtttgcaagaaagtgaagcgtgttgcaactcgcttcgttcgtgaaaaa
gttgttcgtcctatgtgttctagatcccctatgctattatttaagcttaagaaaattatttatgatttgcatttata
tagattaagaaaacagattaggatgttgagacgtcaaaaacagcgcgattatgagttagagtgtgtcactaatctgt

tacaattatcgaatccagtgcaggcaaaaccagagatggataaccctaatccaggacctgatggcgagggtgaagtt

gaattagaaaaggatagcaatgttgttttaacaactcagcgagatcctagtacatctattccagcgccggtgagcgt

aaaatggagtagatggactagtaatgacgtagtagatgattatgccacaatcacatctcgatggtatcagattgctg

aatttgtttggtcgaaggatgatccatttgataaggagttagcacgtttaattttgcctcgtgctttgttatctagt

atagaggctaattctgatgctatatgtgatgtgcctaatactatcccatttaaggtacacgcatattggcgaggcga

tatggaagttagagttcaaattaattcaaataaattccaagttggtcaattacaagctacttggtattattcggatc

atgagaatttgaatatatcgtctaagagaagcgtttatggattttcacaaatggatcatgctttgattagtgcgtca

gcaagtaatgaagcaaaattagttattccatataagcatgtttatccatttttaccgacaagaattgtgccagattg

gactactggcattttagatatgggtgctttgaacattcgtgtaattgctcccttacggatgagtgctactggtccaa

ctacctgtaatgtcgtcgtgtttattaaattaaataacagcgagtttacagggacttcttctggtaagttttatgcg

agccaaatcagggcaaaacctgagatggatcgtatattaaatttggcagagggattgttgaataacacgattggtgg

taataatatggataatccttcttatcaacaatctcctcgtcattttgtcccgactggtatgcacagcttagctttag

gtactaatttagttgaaccattacatgcattacgtttggatgcagccggtacgacacaacatcctgtaggttgtgct

ccggatgaagatatgactgtatcctccattgcatctcgatatggactaattagacgggtacaatggaagaaagatca

tgctaaaggatcacttttgttacaattagatgctgatccatttgtggagcaaagaattgaaggtacgaatccaatat

ctttgtattggttcgcacccgtgggtgtagtatctagtatgtttatgcaatggcgcggttcattagaatataggttt

gatattatagcatcccaatttcatactggtaggttaattgtaggttatgtgcccggtttgacagcatctttgcaact

tcaaatggactatatgaaattgaagtcatcgagttatgtagtatttgatttacaagaaagtaatagcttcactttg

aggtgccatatgtttcatatagaccatggtgggtgcgtaaatatggtggcaattatttaccctcgtcaactgacgct

cctagtacattatttatgtatgtgcaggttccgttgatacctatggaagctgtttcagatactattgatatcaatgt

gtacgtacggggcggtagttcatttgaagtttgtgttccagtccaacctagtttaggtttgaattggaatacagact

ttattttacgtaatgacgaagaatacagggctaagacaggttatgcaccatattatgctggagtgtggcatagcttc

aataatagtaattctcttgtttttaggtggggatctgcttccgatcaaattgctcagtggccgacaatttcagtacc

aagaggtgagctagctttcttacgaattaaggatggaaagcaagctgctgtaggaactcaaccttggcgtacgatgg

ttgtttggccttctggtcatggttataatattggtatacctacgtataatgctgaacgagctcgccagcttgcacaa

cacttatatggtggtggatcattaactgatgagaaggccaaacaattatttgttcctgctaatcaacaaggacctgg

taaggtaagtaatggaaatccggtatgggaagtcatgcgtgcaccattggcaacacagcgtgcgcatattcaagatt

ttgaatttattgaagctattccagaaggagaggagtctcgtaatactacagtcttggatacgaccactactttacag

tcgagtggatttggtcgcgccttctttggagaagcttttaatgaccttaaaacgttaatgcgacgatatcaattata

tggtcaattattattgtccgttactacggataaggatattgatcattgtatgtttaccttcccttgtttaccacaag
ggttagcgttagacattggttctgctggctctccacatgaaatctttaatagatgtcgtgatggtattataccatta
attgcatctggatatagattttatagaggagatttgcgttataagattgtttttccaagtaatgttaatagcaacat
ttgggtacaacatcgaccggatcgtagactggaaggatggtccgcggctaagattgtaaattgtgatgctgtgtcta
ctggtcaaggggtgtataatcatggttatgctagtcacattcaaatcacgcgtgtaaataatgttatagaattggaa
gttccattttataatgctacttgttataattatttacaggcgtttaatgcgtctagcgctgcatctagttatgcagt
atctttaggagaaatatcggttggttttcaagctacaagtgatgatattgcatctattgttaacaaacctgttacta
tttattatagtattggagatggtatgcaattttctcagtgggttggatatcaaccgatgatgatcctagatcagctt
cctgcaccagtagtaagggccgtgcctgagggccctattgcgaagattaaaaacttcttccatcaaacagccgacga
agttagagaagctcaggcagcaaagatgcgtgaagatatgggtatggttgtccaagatgttattggagaacttagcc
aggccataccggatcttcaacaaccggaggttcaagcaaatgtcttctcactggtgtctcagttagtgcatgctatt
ataggtactagtttgaagacagttgcttgggcgattgtttcgatttttgtgaccctaggtttgattggacgtgaaat
gatgcattcagtcataactgtagttaagcggttattagaaaaatatcacttggcgacgcaaccccaggaatccgcca
attcaggtacggttatttccgctgttccagaagcacctaatgctgaagcagaggaggccagtgcctgggtatccatt
atttataatggtgtgtgtaatatgttgaatgtagccgctcaaaaaccgaaacaatttaaagattgggtaaaattagc
taccgtagattttagtaataattgtagaggtagtaaccaggtatttgtatttttcaagaatacatttgaagtgttga
agaaaatgtggggttatgtattttgtcaaagtaatcctgcagcgcgtttgttgaaagctgtgaatgacgagcctgag
attttgaaagcatgggtgaaggagtgtctgtatttggatgatcctaaattcagaatgcgtcgagcgcatgatcaaga
gtatatcgagagagtgtttgcggcacattcatatggacaaattttgctacatgatttaactgctgaaatgaatcaat
cacgaaatttgagtgtgtttacacgtgtgtatgatcaaatttcaaaattgaaaaccgatcttatggaaatgggatca
aatccatatataaggcgtgaatgttttacgatatgcatgtgtggtgcatctggaattggaaaatcatatttgaccga
ttctttatgcagcgagctcttacgtgcgagtcgtactcctgtgacaacaggcataaaatgtgttgttaatccattat
ctgattattgggatcaatgtgattttcagcctgttttgtgcgttgacgatatgtggagtgttgaaacatctactacg
ctcgataagcagttgaatatgcttttccaggttcattctcctatcgtactttctcctcctaaagctgatttagaagg
taagaaaatgcgatataacccggaaatattcatatacaatacgaataaacctttcccgaggtttgaccgtattgcta
tggaagctatttatcggcgtagaaatgttttgattgaatgtaaagcgagtgaagagaagaagcgaggatgtaagcat
tgtgagaatgatattcctattgctgaatgtagtcctaagatgttgcaagattttcatcatatcaaatttaggtatgc
acatgatgtatgtaattctgagaccacatggtctgaatggatgacgtatagtgaatttcttgaatggataactcctg
tgtatatggctaatcgtcgtaaggcgaatgaatcgtttaagatgcgtgtggatgaaatgcaaatgttacgtatggat

gaaccattagaaggtgataatattctcaataagtatgttgaagttaatcagcgcttagtggaggaaatgaaggcatt
taaggaacgcacactatggtcagatttacatcgcgtaggtgcggaaattagtgcgtcagttaagaaagctttaccaa
ccatttccataaccgaaaaactaccacattggactgttcaatgtggcattgctaaacctgagatggatcatgcttat
gaggttatgagttcgtatgcagctggaatgaatgcggagattgaagcgcatgaacaagtccggcgttcatcagtgga
atgtcaatatgcagagcctcaagcttcaagaaatcctgatgatgaagggccaaccatagatgaagaacttatgggcg
acactgaatttacatcacaggctttagaacgtcttgtggatgaaggttatataactggaaaacagaagaaatatata
gctacgtggtgtagtaagcgtcgtgaacatactgctgactttgatcttgtctggactgataatttgcgtgtgttaag
tgcgtatgcgcatgaacgctcatcttcaactcggctttctacggatgatgtcaagttatataaaacaattagcatgt
tacatcaaaagtatgataccacagagtgtgctaaatgtcaacattggtatgctccgttgactgatatctatgttgat
gataagaaattgttttggtgtcagaaagagaaaaagacacttatcgatgtccgcaaattgtcgaaagaagatgtgac
tgttcaatcaaaattgattaatttatctgttccttgtggtgaagtgtgtatgttacattcaaaatatttcaattatc
ttttccacaaagcatggttgtttgagaacccaacttggcgcctaatatataatggtaccaagaagggtatgcctgag
tactttatgaattgtgtggatgaaatttcattagattccaaatttggtaaagtgaaagtatggttgcaagcgatcat
tgataagtatttaactcgtcccgtgaaaatgattcgtgattttcttttcaagtggtggccgcaagttgcgtatgtgt
tgagcttgctaggtataattggtataactgcgtatgaaatgagaaatccgaaaccaacttctgaggaattagctgat
cattatgtgaataggcattgtagctctgattttggtcaccaggactggcatcacctcaaggattgaaatatagtga
agcagtaacagcaaaggcacctagaatccatagattgccagtgactactaagcctcagggatcaactcaacaagtag
acgctgctgtgaataaaattttacagaacatggtttacattggtgttgttttcccgaaagtgcctggtagtaagtgg
cgagatattaattttaggtgtcttatgcttcataataggcaatgtttaatgttgaggcattatattgagtcaactgc
cgcctttcctgaggggaccaagtactattttaagtatattcataatcaagagactagaatgtctggtgatatttctg
gtattgaaattgatttgttgaatttacctagattgtattatggtggtctcgcgggagaggagtcatttgatagcaat
attgtgcttgtgactatgcctaatcgtattcctgagtgtaagagcattattaaatttattgcgtcacataatgaaca
tatacgtgctcagaatgatggagtgttagtaactggcgaccatactcagctattggctttcgagaataataataaga
ctccaataagtatcaacgctgatggtttgtatgaggttatacttcaaggagtatatacttatccataccatggcgat
ggtgtttgtggttccatattgctgtctcggaatttacaacggccgattataggtatccatgttgctggtactgaagg
attgcatggctttggagtcgctgaaccactggtacatgaaatgttcaccggtaaagcaatcgagagtgaaagagagc
cgtatgatcgtgtgtatgaacttccgttgcgtgaattagatgaatctgatattggtttagatactgatttatatccg
attggtagagtggatgcaaagttagctcatgctcaaagcccttctactgggatcaaaaagacgcttatccatggaac
atttgatgtaaggactgaaccaaatccgatgtcgtcacgtgatccaagaatagcgccgcatgatcctttgaagttag

ggtgtgaaaagcatggtatgccttgttcaccgtttaataggaaacatctggaattagcgacaaatcatttgaaagaa
aaattagtttcagtagttaaaccaataaatggttgcaagattagaagtttgcaagatgctgtatgtggtgtgcctgg
tttagatgggtttgattcgatatcttggaatactagtgctggttttcctttgtcttcattaaagccacctggaacat
ccggtaagcgatggttgtttgacattgagctgcaagactcgggatgttatctcctgcgtggaatgcgtcccgaactt
gagattcaattatcaacgacacagttaatgaggaaaaaggaataaaacctcacactatattcacggattgtttgaa
agatacttgtttgcctgttgaaaaatgtagaatacctggtaagactagaatatttagcataagtccggtgcagttta
ccataccgtttcgacagtattatttagactttatggcatcctatcgagctgcacgacttaatgctgagcatggtatt
ggtattgatgttaacagcttagagtggacaaatttggcaacaaggttgtctaagtatggcactcacatcgtgacagg
agactataagaattttggtcctgggttagattccgatgttgcagcttcagcgttcgaaattattatcgactgggtat
tacattacactgaagaagataataaagacgaaatgaagcgagtaatgtggaccatggcgcaagagatcttagcgcct
agtcatctatgtcgcgacttggtgtaccgagtaccttgcggaattccatcaggttctccaataacggacatattgaa
tacaatttcaaattgtttgttaattaggttagcctggttaggtattactgatttgcctttgtccgagttctctcaaa
atgttgttcttgtttgttatggtgatgatcttatcatgaatgttagtgataacatgattgataaatttaatgctgtg
acaatagggaaattcttttcacaatataagatggaatttacggatcaggacaaatcaggaaatactgtgaagtggcg
gacgttacagactgctactttcttgaagcatgggtttttaaaacatccaactagacctgtgtttctggctaacctag
acaaggtttcggtagaaggaacgacgaattggacccatgctcgaggattgggtcgtcgtacagcaaccatagagaat
gctaagcaagcgctagagttagcattcggatggggtccagaatactttaactatgtcagaaatactattaaaatggc
ttttgacaagttgggtatttatgaagaccttatcacatgggaagaaatggatgttagatgttatgctagcgcgtagt
atttaattttgaatacttattagttttaattttattttaggttattggaattgagggaagtaccacccccccaagacc
ttcgttttaaatctactaaaaggagtgaacctatatataagagtctaacgacagagtggatcagaccaccatcttta
gcttatatatgggaaaggttgagttgcctctaaagactcagctccatagtagagtagttttaattacgattaaagtg
gtactctaggttaggtgttactcgcgtattatcaactagtggtaatgcgtcctaattttagtatagttttaaccata
atagtaaaaaaaaaaaaaaaaaaaaaaaaa

>VVV-mCherry ($VVV_{mC}$)

tttaaaattcgctatgggaggcgatttatgccttccatagcgaattacggtgcaactaacaattttagata
gtagccatgaacaaacattatgattactcactacgtattgatcatttttttcaatggcttgcgtagcatgaagcgcat
gcttgtagttataactatgttattttgcaagttggagataattgtattggattatggatgcgtgcactaagtgtcta
catctatagtcgtttgtggttcaagtttttgtgttagtagtacaatcttgaagaatgtaagtatcgtatgaatgata
tttgaatgacaacactgaagtataaaatatataaaatccaaaaatatttttaatcttattcagtgtagtgtttgata

gagtagaatgccatgtgaccgctcaaagaagtccattatggtatatcattcgaagtcgaatacttgtgtatagttat
tgtattttattagtaatattagtagtccgtaactatcataatcctattatagtttgattatatgatagaccactgca
gtatcgagtagagtttagaaagagtagtgcaatagtaagatcactgtcaccgaccactcattgtaatagtgaggttt
gtcggaaaccagttattgtgcagcgactagcaatcgtgaatcaatatagttggtattctaaatatgagacgattcgg
cgattttattgcgactgaaatttcatatttagcatgtcaggtcttattatgaatgctcgagtatttatttctgcggt
agagtagggacccctctatctctcaggtactgtatgaggcgaaagtgtgaaagtaacttatgtctctatacataagt
gactgtatcgggatttcctttggcaagaatccttttaatacagtataatttatgctacggtacgttacgttcgcagg
gcacccgttaatgtcacatagcccagacgatgacgaatggaaagacattacttttattttaatgctacgattattg
ctgttttattttgctgtttttatttgctattatattttgctattttcattattgctaaatatatttctttgctattt
ttgctttatatattagattcaattcttttattttatattttcaatttgattttgattttgaaggtaaatatatata
aaaatggcatttagttgtggaactctttcttatgctgctgttgcccaagctccctctgtagctcatgctccccgtag
ttgggagattgatgaagctaggcgtcgacgcgttatcaagcgtttggcgttggaacaggaacggattcgaaacgttc
tcgacgtcactgtgtatgatcatacaacgtgggagcaagaggatgcgcgtgataatgagttccttacggaacaattg
aataatttatatacgatatattctatagctgaaagatgtacccgccggcctgttcaagaacatgtccccatttcaat
cagtaatagatattcccctttagaatcccttaagattgaggtaggaaaagacgcgggtgagttcgtatttaagaaac
ccaaatatacaaagatttgtaagaaagtgaaacgggtggcatcaaaatttgtgcgcgagaaagttgttaggcccgtt
tgtaatcgatcgccaatgttattatttaaaattaagaaagtaatatatgatttacatttgtatcggttacggaaaca
agttcggcttctcagacgcgaaaaacagcgtgaatatgagttagagtgtgttactagtttgctacagctatctaatc
ctgtttcagctaaacctgag**atggacaatcctatggtgagcaagggcgaggaggataacatggccatcatcaaggag**
**ttcatgcgcttcaaggtgcacatggagggctccgtgaacggccacgagttcgagatcgagggcgagggcgagggccg**
**cccctacgagggcacccagaccgccaagctgaaggtgaccaagggtggccccctgcccttcgcctgggacatcctgt**
**cccctcagttcatgtacggctccaaggcctacgtgaagcaccccgccgacatccccgactacttgaagctgtccttc**
**cccgagggcttcaagtgggagcgcgtgatgaacttcgaggacggcggcgtggtgaccgtgacccaggactcctccct**
**gcaggacggcgagttcatctacaaggtgaagctgcgcggcaccaacttcccctccgacggccccgtaatgcagaaga**
**agaccatgggctgggaggcctcctccgagcggatgtaccccgaggacggcgccctgaagggcgagatcaagcagagg**
**ctgaagctgaaggacggcggccactacgacgctgaggtcaagaccacctacaaggccaagaagcccgtgcagctgcc**
**cggcgcctacaacgtcaacatcaagttggacatcacctcccacaacgaggactacaccatcgtggaacagtacgaac**
**gcgccgagggccgccactccaccggcggcatggatgaattgtacaaagctaaacctgag**atggacaatcctaatcct
ggtccagatggcgaaggtgaagttgaattagaaaaagatagtaatgtagtattaactacacaacgtgatccgagtac

ttctattcctgctccaactagtgtgaagtggagtagatggactagtaatgatgttgtggatgattatgccactataa
cttcgcgttggtatcagattgccgaatttgtatggtcaaaggatgatccatttgataaggaattggcgcgtttaatt
ttacctcgagctttgttatctagtattgaggctaattctgacgctatttgtgatgtacctaatactattccgtttaa
ggtacatgcatattggcgtggagatatggaagttcgagtgcagattaactcgaataaattccaggttggtcaattgc
aggcaacttggtactattcggatcatgaaaatttgaatatccagacgaagcgaagtgtgtatggtttttcgcatatg
gatcatgctttgattagcgcatcagcgagtaatgaagcaaaattaatgatacctttaaacatgtatatccattctt
accaacgcgtgtcgttcctgattggacaactggtattcttgatatgggtaccttaaatattcgtgtaattgctccac
tacgtatgagtgcgacgggaccaaccacttgtaatgttgtagtatttattaagttaaataatagtgaattcactggt
acttcttctggtaagttttacgcgaatcaaatcagggcaaaacctgaaatggaccgtgtgttaaatttggcagaagg
attactaaataatactgtaggtggttgtaatatggataatccgtcatatcagcaatctccgcgtcattttgttccta
ctggtatgcatagtttagctttaggcactaatttagtagagcctttgcatgcattacgattagatgcatcaggtaca
acacaacatccagttgggtgtgcgcctgatgaagatatgactgtatcttccattgcatcacgatatggtttaattcg
ccaagtgcaatggaagaaagaccatgcgaaaggatcattattattacaacttgacgctgatcctttcgttgaacaga
aaattgagggaaccaatccaatttctttgtattggtttgctccagttggagtcgtatctagtatgtttatgcaatgg
agaggttctttagaatatagatttgatattatagcttcccaatttcatacgggtaggttaattgtaggttatgttcc
tggactgacagcttctttacaacgtcaaatggactatatgaaattgaaatcatctagttatgtggtgtttgatttac
aggaaagtaatagttttacgtttgaagtgccctatgtgtcatacagaccgtggtgggtgcgtaagtatggtggtaat
tatctgccatcttctactgatgcgcctagcacactgtttatgtatgtacaagtaccattgatacctatggaagctgt
ttctgatactatagatatcaatgtgtatgtgcgtggtggcagttcgtttgaggtttgtgttccagtccaacctagtt
taggtttgaactggaatacagatttcatattacgtaatgatgaagagtaccgcgcaaagaatggatatgcaccatat
tatgctggtgtgtggcatagcttcaataatagcaattcgcttgtttttagatggggttcggcttcagatcaaattgc
tcaatggccaacaataacagtgcctcgaggagagttggcattcttgcgtatccgcgatgctaagcaagctgctgtag
gaacgcaaccttggcgtactatggtcgtttggccttcaggtcatggatataatattggaataccaacttataatgct
gaacgagcaagacaacttgctcagcatttgtatggtggtgggtctttgacagatgaaaaggctaagcaattatttgt
gcctgctaaccagcaaggacccggcaaagtaagtaatggtaaccctgtctgggaagtaatgcgcgcgcctcttgcaa
ctcagcaagcgcatatacaagattttgaatttgttgaagctgttccagaaggcgaagaatcacgcaacactacggtg
ctagatacgacaataacgttacagtctagcggatttggtcgcgctttcttcggtgaggcatttaacgatcttaagac
gttaatgcgccgataccaattatatggtcaattattgttatccgttactacggataaggatattgatcattgtatgt
ttaccttcccttgtttacctcaagggctagcgttagatataggttcggctggatctcctcatgaaatatttaatcgc

tgccgtgatggtatcattccattgatagcgtcagggtatcggttttatcgaggcgatttacggtttaaaattgtttt

cccaagtaacgttaatagcaatatttgggtacaacaccgaccagatcgtagactgaaaggatggtctgaagcgaaaa

tagtaaactgtgatgctgtatctactggacaaggcgtttataatcatggatatgctagtcatattcagattacgcgt

gtaaataatgttatagaattggaagtcccgttttataacgctacgtgctataattatttgcaagcgtttaacccatc

tagtgcagcgtcaagttatgccgtttcgctcggagagatttcggttggttttcaagctactagcgatgacattgcag

ccatagttaataaacctgtaactatatattacagtattggcgatggtatgcagttttcgcagtgggttggttatcaa

ccaatgatgattctagatcaattgccagcaccagtagttagggctgtgcctgagggccctatagcgaagataaagaa

ctttttccatcaaacggcagatgaagttcgagaagctcaggccgcaaagatgcgtgaagatatgggtatagtagtcc

aagacgttataggagagttaagtcaggctatacccgatcttcaacaaccggaagttcaagcgaatgtttttttctctg

gtgtcacagttagtgcatgctatcatcggtactagtcttaagacagttgcttgggcgattgtttcgatttttgtaac

cctaggtttgattggacgtgaaatgatgcattcagtcataactgtagttaagcggttattagaaaaatatcacttgg

cgacgcaaccccaggaatccgccaattcaggtacggttatttccgctattccagaagcacccaatgctgaagcagag

gaggccagtgcctgggtatccattatttataatggtgtgtgtaatatgttgaatgtagccgctcaaaaaccgaaaca

atttaaagattgggtaaaattagctaccgtagattttagtaataattgtagaggtagtaatcaggtatttgtgtttt

tcaagaatacgtttgaagtgttgaagaaaatgtggggttatgtgttttgtcagagtaatcctgcagcgcgactcttg

aaagcagtgaatgatgaacctgagattttaaaagcgtgggttaaagaatgtctgtatttagatgatcctaaatttag

aatgcgacgtgcgcatgatcaagagtatattgagagagtgtttgcggcccattcgtatggacaaattttattgcatg

acttaacggctgaaatgaatcaatcgcgtaatttaagtgtgtttacgagagtgtatgatcaaatatctaaattgaag

acggatctcatggaaatgggatcaaacccatatatcaggcgtgaatgctttacgatttgtatgtgtggtgcatctgg

aattggtaagtcttatttaactgattctttatgcagcgagctcttacgtgcgagtcgtactccagtgacaacgggca

ttaagtgtgtcgtgaaccctttgtctgattattgggatcagtgtgattttcagcccgttttatgtgttgatgacatg

tggagtgttgaaacgtctactacgctcgataaacagttaaatatgctatttcaggttcattcaccaattgtactttc

acctcctaaagctgatttagaaggtaagaaaatgcgttataatcctgaaatattcatatataatacgaataaacctt

ttccgaggtttgatcgtatagctatggaagctatttatcgacgtagaaacgttttaattgaatgtaaggctaatgaa

gagaagaagcgtggatgtaaacattgtgagaataatatacccattgctgaatgtagtccaaaaattttgaaagattt

tcatcacattaaatttcgttatgctcatgatgtgtgtaattctgaaacgacgtggtctgagtggatgtcgtataatg

aatttttggaatggattactcctgtatatatggctaatcgacgtaaagcaaatgaatcgtttaagatgcgtgttgat

gaaatgcaaatgttgcgtatggatgagcccttggaaggcgataatattttaaataagtatgttgaagttaatcagcg

cttagttgaggaaatgaaagcttttaaagagcgaaccctctgggctgatttacaacgtgttggctcagagattagta cttcagttaagaaagcattaccaactatttccattactgagaagctaccacattggactatccaatgtggcatagct

aagcctgaaatggatcatgcttatgaagttatgagttcatatgcagcaggaatgaacgcagaaattgaagcgcatga

acaagttcgtcgttcttctttggaatgtcagtgtattgagccttcaacttcaagacctctggatgaagagggtccta

ctatcgacgaggaattacttggcgaagtagaatttacttcttcagctttggagcgtttggttgatgaggggtatatt

actggtaaacaaaagaagtacatggcaacttggtgtacgaaacgaagagagcatgtatccgattttgatttagtatg

gacggataatttgcgtgttttgagtgcgtatgtccacgagcgttctacatctacgcgtttatctaccgatgatgtta

aattatttaagacgattagtatgttacatcagaggtatgataccactgattgtgcaaaatgccaacattggtatgca

ccattaacagctatttatgttgatgatagaaagctattttggtgccagaaggagactaagactttgatagatgttcg

taaattgtcgaaagaggacgttacagtccaatcgaaattaattaacttatcggttccgtgcggtgatgtatgtatgt

tacattctaagtactttaattatttattccataaagcgtggttgtttgaaaatccaacatggcgtttaatatataat

ggtactaagaaaggtatgcctgagtatttcatgaattgcgtggatgaaatttcattagattcaaaattttgtaaagt

aaaggtttggcttcaagcaattattgataaatatttgactcgtccagtgaaaatgattcgtgactttctatttaaat

ggtggccgcaagtagcatacgtgttaagtttgttaggtataattggtataactgcgtatgagatgcgtaatcctaaa

tcaacagcagaagacttggctgagcactatgttaataggcattgtagttcagattttggtcaccaggtatggcgac

tcctcagggattaaaatatagtgaagcgataacagctaaagcgcctagaatccatagattgcccgttactactagac

ctcagggatcaacgcaacaagttgacgccgctgtgaataagattttgcagaatatggtgtatatcggtgttgtgttt

ccgaaagtgcctggtagtaagtggcgagatattaatttagatgtcttatgcttcataatcggcaatgtttgatgtt

gcggcattacattgagtcgacggctgcttttccggagggtaccaaatactattttaagtatattcataatcaagaaa

ctcgaatgtcaggtgatatatctggtattgagattgatttattgagtttacctagattgtattatggtggcttagct

ggggaagagtcgttcgatagcaatatagtgttagtaactatgccgaatagaattcctgagtgtaagagtattgtgaa

gtttatagcttcacatgctgaacatgctcgtgctcaaaatgatggtgtgttagttactggtgaacatactcagttat

tggcattcgagaataataataaaacacctataagtattaatgctgatggtttgtatgaggttatacttcaaggagta

tacacttatccataccatggtgatggtgtttgtgggtctatattattgtctcgtaatttacaacgaccgattatagg

gatccatgtagctggtactgaaggattacatggctttggtgttgctgaacctcttgttcatgagatgttcactggga

aagcaatagagagtgaaagggaaccgtatgatcgtgtgtatgaattacctttgcgtgaattagatgaatctgatata

ggtttagatactgatttatatcctataggaagagttgatgcgaaattagctcatgcccaaagtccttcaacaggaat

taaaaagacgcttattcatggtacttttgatgttcggactgaaccgaatccgatgtcatcacgagacccaagaatag

caccacatgatccgttgaagttagggtgtgagaaacatggtatgccatgttctccatttaatcgaaaacatttggaa

ttagcaacgactcatttaaaggagaagttaatttccgtagttaaacctataaacggatgcaagattagaagtttgca

agatgctgtgtgtggtgtaccaggtttggatggctttgattcaatatcctggaatactagtgctggttttcctttat
cttcattaaaaccgccaggctcttctggtaagcgatggttgtttgatattgaattacaagattcaggatgttatctt
ttgagagggatgagacctgaacttgagatacagttgacaacaactcagttaatgaggaagaagggaataaagcctca
cactatattcacggattgtttgaaagatacatgtttgcctgtggaaaaatgcagaatacctggtaagactagaatat
ttagtataagtcccgtccaatttacgattccattccgacaatactatctcgattttatggcgtcgtaccgtgccgct
agacttaatgctgagcatggaataggtatagacgtgaacagcttggaatggacaaacttggcaacaagtctgtcgaa
gtatggcacgcatattgtgacaggagattacaagaattttggtcctgggttagattctgatgttgccgcttcagctt
tcgaaattatcattgattgggtgttaaattacactgaagaagatgataaagacgaaatgaagcgtgtaatgtggact
atggctcaggaaatcttagctcctagtcacttatgtcgtgatttagtatatcgcgtaccatgcggtattccttctgg
atcaccaattacggacattttgaatactatttcgaattgtttgttaattcgattggcttggcaaggtattactgatt
tgcctttatccgaattttctagacatgtcgtgctagtttgttacggtgatgatctcatcatgaatgtaagtgatgag
atgatagataaattcaacgctgtaacaattggcgatttcttttcgcgatataagatggaatttacggatcaggataa
atctggaaatacagtgcggtggcgaactttacaaactgccacgtttttgaagcatgggttcttgaaacatccaacaa
gacccgtgtttctagccaatctggataaggtttctatagaaggaacaaccaattggacacatgctcgaggattgggt
cgtcgagtagcaaccattgagaatgctaaacaagcgctagagttggcattcggatggggtcccgaatactttaatca
tgttcggaataccattaaaatggcattcgacaagttaggtatttatgaagatcttatcacatgggaagaaatggatg
ttagatgttatgctagcgcgtagtatttaattttgaatacttattagttttaattttattttaggttattggaattg
agggaagtaccacccccaagaccttcgttttaaatctactaaaaggagtgaacctatatataagagtctaacgaca
gagtggatcagaccaccatctttagcttatatatgggaaaggttgagttgcctctaaagactcagctccatagtaga
gtagttttaattacgattaaagtggtactctaggttaggtgttactcgcgtattatcaactagtggtaatgcgtcct
aattttagtatagttttaaccataatagtaaaaaaaaaaaaaaaaaaaaaaaaaaa
